# Agent-based model demonstrates the impact of nonlinear, complex interactions between cytokines on muscle regeneration

**DOI:** 10.1101/2023.08.14.553247

**Authors:** Megan Haase, Tien Comlekoglu, Alexa Petrucciani, Shayn M. Peirce, Silvia S. Blemker

## Abstract

Muscle regeneration is a complex process due to dynamic and multiscale biochemical and cellular interactions, making it difficult to identify microenvironmental conditions that are beneficial to muscle recovery from injury using experimental approaches alone. To understand the degree to which individual cellular behaviors impact endogenous mechanisms of muscle recovery, we developed an agent-based model (ABM) using the Cellular Potts framework to simulate the dynamic microenvironment of a cross-section of murine skeletal muscle tissue. We referenced more than 100 published studies to define over 100 parameters and rules that dictate the behavior of muscle fibers, satellite stem cells (SSC), fibroblasts, neutrophils, macrophages, microvessels, and lymphatic vessels, as well as their interactions with each other and the microenvironment. We utilized parameter density estimation to calibrate the model to temporal biological datasets describing cross-sectional area (CSA) recovery, SSC, and fibroblast cell counts at multiple time points following injury. The calibrated model was validated by comparison of other model outputs (macrophage, neutrophil, and capillaries counts) to experimental observations. Predictions for eight model perturbations that varied cell or cytokine input conditions were compared to published experimental studies to validate model predictive capabilities. We used Latin hypercube sampling and partial rank correlation coefficient to identify *in silico* perturbations of cytokine diffusion coefficients and decay rates to enhance CSA recovery. This analysis suggests that combined alterations of specific cytokine decay and diffusion parameters result in greater fibroblast and SSC proliferation compared to individual perturbations with a 13% increase in CSA recovery compared to unaltered regeneration at 28 days. These results enable guided development of therapeutic strategies that similarly alter muscle physiology (i.e. converting ECM-bound cytokines into freely diffusible forms as studied in cancer therapeutics or delivery of exogenous cytokines) during regeneration to enhance muscle recovery after injury.

## Introduction

Skeletal muscle injuries account for more than 30% of all injuries and are one of the most common complaints in orthopedics^1,2,3^. The standard treatment for muscle injuries is limited mostly to rest, ice, compression, elevation, anti-inflammatory drugs, and immobilization^1^. These treatments lack a firm scientific basis and have varied outcomes, some resulting in incomplete functional recovery, formation of scar tissue, and high injury recurrence rates^4,5^. Our fundamental understanding of the individual cellular and subcellular behaviors of muscle cells has advanced and made it clear that interactions between cells and their microenvironment is critical for healthy regeneration. These interactions are dynamic, involve feedback mechanisms, and lead to complex emergent phenomena; therefore, there are numerous possible interventions that could enhance muscle regeneration.

Muscle regeneration requires an abundance of cells and cytokines to interact in a highly coordinated mechanism involving five interrelated cascading phases including: degeneration, inflammation, regeneration, remodeling, and functional recovery^6^. Following an acute muscle injury, there is a time-dependent recruitment of neutrophils, monocytes, and macrophages to remove necrotic tissue and release factors that regulate fibroblast behavior and SSC activation, proliferation, and division^7^. Following initial inflammatory response, fibroblasts and SSCs activate and proliferate with the macrophages shifting from their pro to anti-inflammatory phenotype. In healthy muscle, this process would be followed by remodeling of the muscle where the fibroblasts apoptose and SSCs differentiate and fuse to repair the myofibers^8^. Each cell involved in this process secretes cytokines that help regulate cell recruitment and chemotaxes to modulate the dynamics of the recovery. It has also been shown that the molecular events implicated in angiogenesis occur at early stages of muscle regeneration to restore microvascular networks that are crucial for successful muscle recovery^9^.

There are numerous cytokines involved in muscle regeneration, many of which have been individually studied to examine their influence on muscle regeneration^10^. These cytokines play key roles in dictating cell behaviors and are major drivers of the regeneration cascade^11^. The dynamics of these cytokines control many aspects of the microenvironment and altering their properties to optimize treatments has been proposed in a variety of settings^12^. Testing alterations in cytokine dynamics experimentally has proven to be complex and expensive due to difficulties in cytokine identification and quantification as well as confounding factors due to pleiotropic activities of cytokines and interactions with soluble receptors^13^. These challenges make it difficult to holistically test different diffusion and decay properties for numerous cytokines^14^. However, if we could better understand the synergistic effects of alteration in cytokines, we could design a more effective therapy for treating muscle injury.

There are over a million possible combinations of cytokine alterations, making it unrealistic to study all combinations with experiments alone. For this reason, an *in silico* approach is needed to fully explore the possible treatment landscape and make predictions on potential targets to enhance muscle recovery. Over the last several years, Agent Based Models (ABM) of muscle regeneration have developed to study muscle regeneration in a variety of applications^8,15–19^. These models were foundational for exploring the role of SSCs in a variety of muscle milieus^8,15,17^ and for demonstrating how ABMs can be used to simulate therapeutic interventions^16^. However, previous models employed simplistic, non-spatial representations of cytokine behaviors and properties, which limited their ability to recapitulate cytokine alterations such as injection of TGF-β^19^. Furthermore, these prior models did not include microvessel adaptations and dynamic ECM properties which are crucial for understanding the altered microenvironmental state following muscle injury. These critical limitations must be addressed in order for ABMs of muscle regeneration to provide meaningful insights into treatments for muscle injury.

The goals of this work were to: 1) develop an ABM of muscle regeneration that includes cellular and cytokine spatial dynamics as well as the microvascular environment, 2) calibrate the model to capture cell behaviors from published experimental studies, 3) validate model outcomes by comparison with multiple published experimental studies, 4) conduct *in silico* experiments to predict how altering cytokine dynamics impacts muscle regeneration. For model calibration, we implemented an iterative and robust parameter density estimation protocol to refine the parameter space and calibrate to temporal biological datasets^20^. Partial rank correlation coefficient (PRCC) was used to guide *in silico* experiments by identifying parameters and timepoints that were most critical for ideal regeneration metrics.

## Methods

### Agent-Based Model Development Overview

ABMs represent the behaviors and interactions of autonomous agents, such as cells, which are governed by literature-derived rules^15,21,22^. Agent-based modeling provides an excellent platform for studying complex cellular dynamics because they reveal how the interactions between individual cellular behaviors lead to emergent behaviors in the whole system.

We implemented the ABM in CompuCell3D (version 4.3.1), a Python-based modeling software^23^. The ABM’s code is available for download (https://zenodo.org/records/10403014). To build the model, we extended upon about 40 rules developed in previous agent based models of muscle regeneration^8,15,16^ in combination with a deep literature search referencing over 100 published studies to define approximately 100 total rules that dictate the behavior of fiber cells, SSC, fibroblasts, neutrophils, and macrophages, as well as their interactions with the microenvironment, including microvasculature remodeling and cytokine diffusion and secretion (Fig. 1). For a rule to be incorporated into the model, there had to be an established understanding within the literature supporting the behavior (i.e. multiple studies reporting similar findings or supported by other reputable publications). When available, we used experimental data to define the parameters associated with the model rules. There were 52 parameters that could not be related to known physiological measurement; therefore, these parameters were calibrated using parameter density estimation which will be described below in *Model Calibration*. Following calibration of model parameters, separate model outputs were validated by comparison with experimental data, and various model perturbations were conducted and compared to literature results. This process allowed us to have confidence in the predictive capabilities of the model so that we could simulate and predict the sensitivity of muscle regeneration to changes in cytokines.

**Figure 1.**
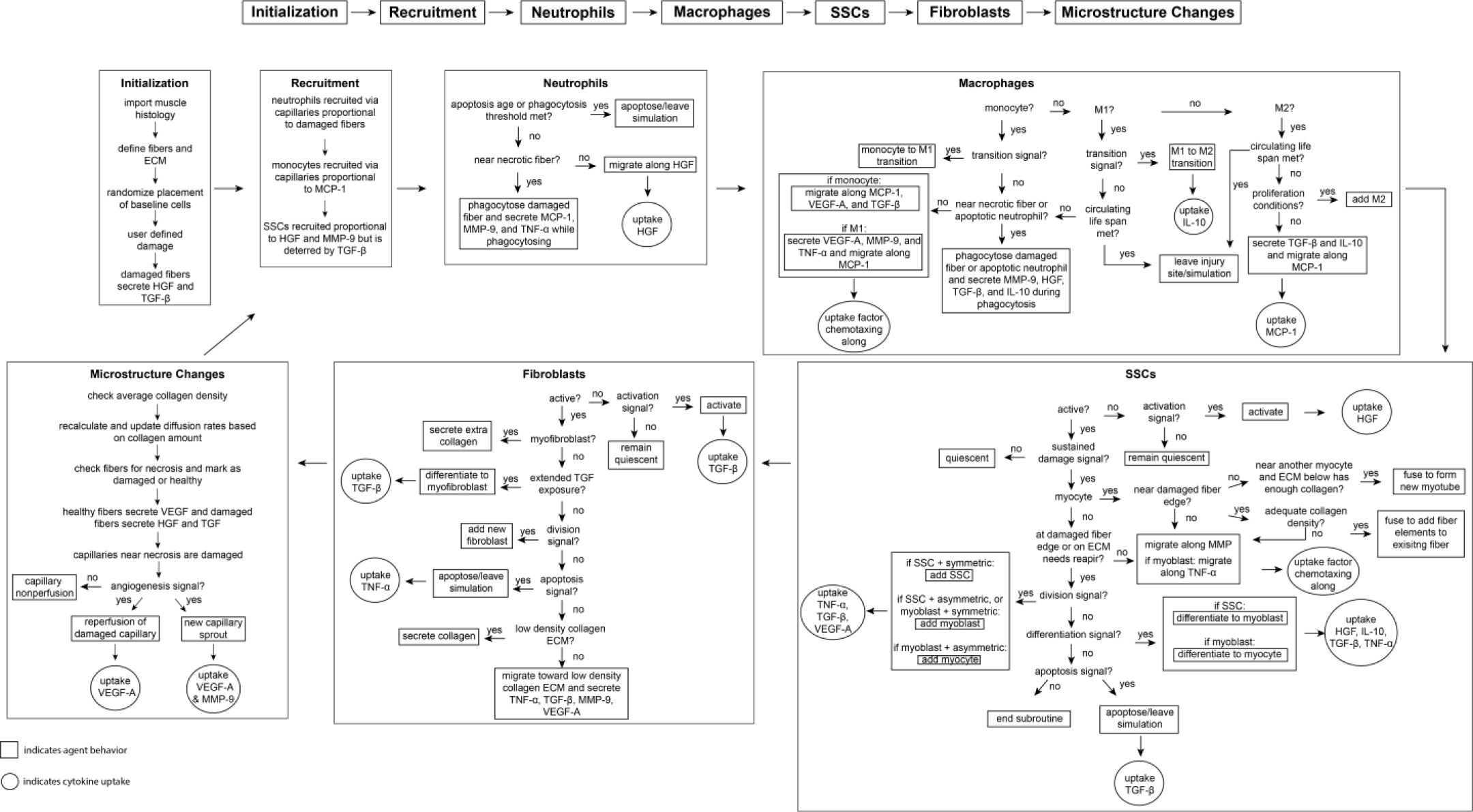
Flowchart of ABM rules. The model starts with initialization of the geometry and the prescribed injury. This is followed by recruitment of cells based on relative cytokine amounts within the microenvironment. The inflammatory cells, SSCs, and fibroblasts follow their literature defined rules and probability-based decision tree to govern their behaviors. The boxes represent the behavior that the agent completes during that timestep given the appropriate conditions and the circles represent the uptake that occurs as a result of the simulated binding with microenvironmental factors for certain cell behaviors. ABM, agent-based model; SSC, satellite stem cell; ECM, extracellular matrix; TGF-β, transforming growth factor β; HGF, hepatocyte growth factor; TNF-α, tumor necrosis factor α; VEGF-A, vascular endothelial growth factor A; MMP-9, matrix metalloproteinase-9; MCP-1, monocyte chemoattractant protein-1; IL-10; interleukin 10.

### Cellular-Potts Modeling Framework

Prior work to construct computational models to represent muscle recovery have used ordinary differential equations^24^ or agent-based modeling software, such as software such as Netlogo^16^ or Repast^19^. While these models have yielded great insights into skeletal muscle damage and recovery processes, they have limited capacity to represent the spatial diffusion of cytokines accurately and explicitly throughout the skeletal muscle. The Cellular-Potts model framework^23^ (CPM, also known as the Glazier-Graner-Hogeweg model), proved an ideal choice because it allows for logic-based representation of cellular behavior and interactions characteristic of agent-based modeling (see Supplemental Text 1 for additional details on CPM).

### ABM Design

The ABM spatially represents a two-dimensional male murine skeletal muscle fascicle cross-section of approximately 50 muscle fibers (Fig. 2A). The ABM depicts the microenvironment of the cross-section as well as the spatial migration of cells and diffusion of various cytokines. The ABM simulates the emergent phenomenon of muscle tissue from an acute injury over the course of 28 days. The spatial agents in the model include muscle fibers, necrotic muscle tissues, extracellular matrix (ECM), capillaries, lymphatic vessels, quiescent and activated fibroblasts, myofibroblasts, quiescent and activated SSCs, myoblasts, myocytes, immature myotubes, neutrophils, monocytes, resident macrophages, pro-inflammatory macrophages (M1), and anti-inflammatory macrophages (M2). In addition, the ABM includes seven diffusing factors, such as hepatocyte growth factor (HGF), monocyte chemoattractant protein-1 (MCP-1), matrix metalloproteinase-9 (MMP-9), transforming growth factor beta (TGF-β), tumor necrosis factor-alpha (TNF-α), vascular endothelial growth factor A (VEGF-A), and interleukin 10 (IL-10). A review of the literature led us to determine that these factors and cytokine isoforms were most critical for representing the behaviors of each cell during the regeneration cascade^25,26^.

**Figure 2.**
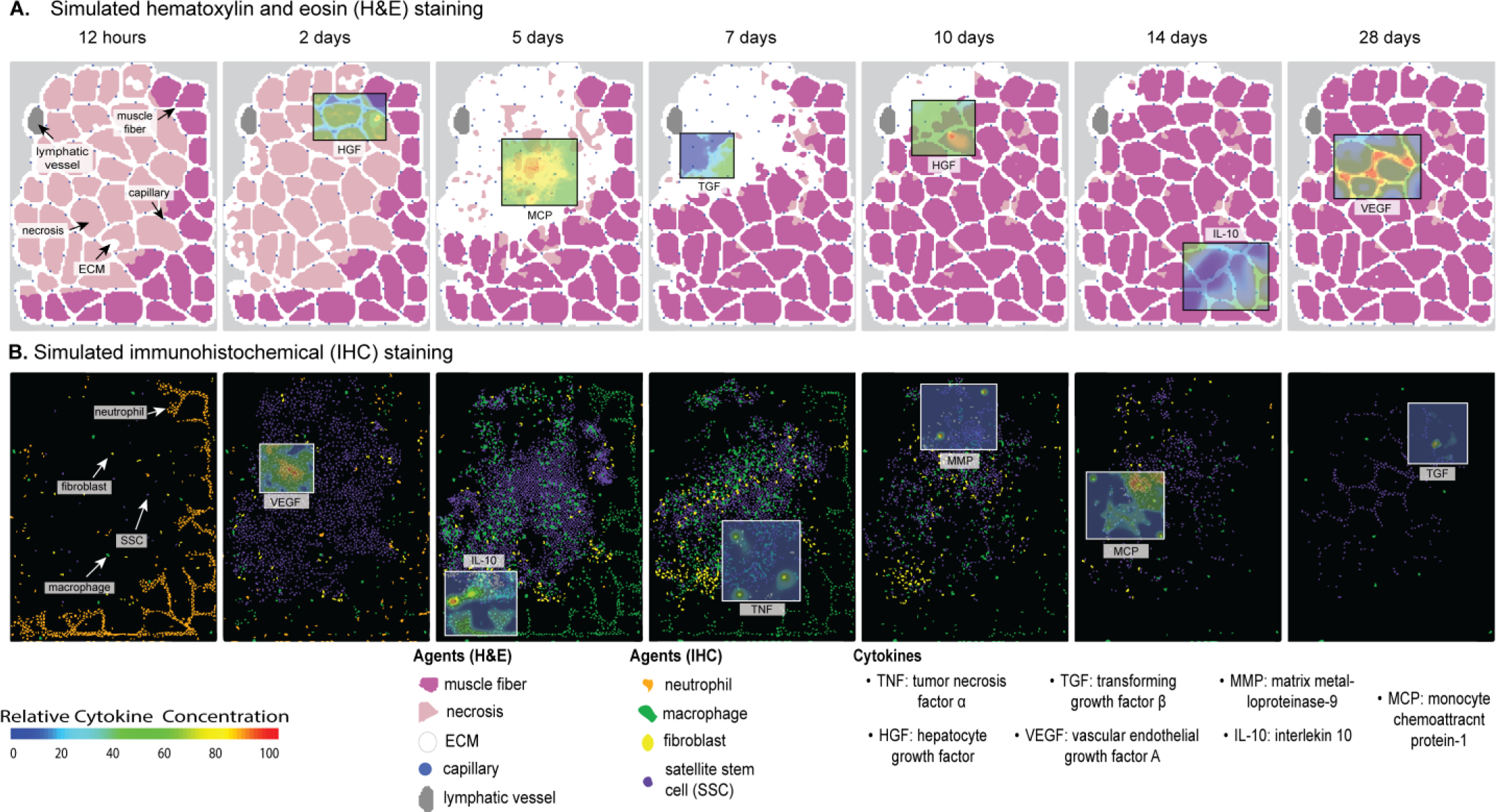
Overview of ABM simulation of muscle regeneration following an acute injury. **A.** Simulated cross-sections of a muscle fascicle that was initially defined by spatial geometry from a histology image. Muscle injury was simulated by replacing a section of the healthy fibers with necrotic elements. In response to the injury, a variety of factors are secreted in the microenvironment which impacts the behavior of the cells. The colors correspond with those typically seen in H&E staining. **B.** ABM screen captures show the spatial locations of the cells throughout the 28-day simulation. The agent colors were matched to those typically seen in IHC-stained muscle sections.

The muscle cross-section geometry was created by importing a histology image stained with laminin α2 into a custom MATLAB script that masked the histology image to distinguish between the fibers and ECM. The mask was imported into an initialization CC3D script that defined the muscle fibers, ECM, and microvasculature to specific cell types and generated a PIF file that was imported into the ABM as the starting cross-section. The injury is simulated by stochastically selecting a region within the cross-section to replace the fiber elements with necrotic elements, where the percentage of CSA damage is an input parameter. When a threshold of fiber elements within a muscle fiber becomes damaged the entire muscle fiber turns necrotic and requires clearance. If the damage is below the threshold, only the region of necrosis must be removed and the SSCs can fuse to the remaining fiber. During model initialization, the injury criteria can be altered to simulate various degrees of myotoxin injury by changing the percent of necrotic tissue following injury.

Each Monte Carlo step (mcs) represents a 15-minute timestep, and the model simulations were run until 28 days post-injury. The cell velocity is limited by how many times the Cellular-Potts algorithm is run, so we set 45 Cellular-Potts evaluations per mcs to ensure stability in migratory agent behavior. The number of Cellular-Potts evaluations per mcs and the lambda chemotaxis parameters were tuned in a simplified simulation of individual cells and their respective chemotactic gradients so we could obtain cell speeds that were consistent with speeds derived from literature sources (Table 7). At each mcs, the agent behaviors are governed by rules that were derived from experimental data found in the literature. The behaviors of each agent are based on environmental conditions, such as nearby cells and cytokine gradients, as well as probability-based rules. As an example, a capillary located near a damaged fiber has a probability of becoming non-perfused and then senses the amount of VEGF-A and MMP-9 at its location to decide if the levels are adequate to induce angiogenesis (Table 6). Model outputs include CSA recovery (sum of total healthy fiber elements normalized by the initial CSA), capillary and collagen density, cell counts, relative cytokine abundance, and spatial coordinates of cells and cytokines.

### Overview of Agent Behaviors

Simulated behaviors (Fig. 2B) of the neutrophils and macrophages include cytokine-dependent recruitment, chemotaxis, phagocytosis of damaged fibers (neutrophils, monocytes, and M1 macrophages), phagocytosis of apoptotic neutrophils (monocytes and M1 macrophages), secretion and uptake of cytokines, and apoptosis. The SSC and fibroblast agent behaviors also include cytokine-dependent recruitment, chemotaxis, secretion and uptake of cytokines, and apoptosis, in addition to quiescence, activation, division, and differentiation. The biological intricacy of some cell types, such as SSCs which have a more complex cell cycle and are regulated by dynamic interplay of intrinsic factors and an array of microenvironmental stimuli, led to the necessity for adding more rules that govern their behaviors^27^. The SSCs have 33 parameters dictating the 17 agent rules (Table 3), fibroblasts have 27 parameters for 11 agent rules (Table 4), macrophages have 31 parameters for 15 agent rules (Table 2), neutrophils have 18 parameters for 7 agent rules (Table 1), fibers have 18 parameters for 4 agent rules (Table 5), and microvessels have 22 parameters for 6 agent rules (Table 6). At each mcs, cytokines are secreted by agents if certain conditions were met. For cell recruitment, the levels of recruiting cytokines for each agent are checked, and if the concentration is high enough to signal cell recruitment (Supplemental Table 1), a new agent is added to the field at the location of the highest concentration. The agents also undergo chemotaxis by sensing the surrounding cytokine gradients and move towards higher concentrations of cytokines, binding and removing that cytokine as they move along it to simulate physical binding of the cytokine to the receptor. Agents that are in a quiescent state require a certain threshold level of cytokines to become activated and cannot chemotax, secrete, divide, or differentiate until this threshold is reached. Our model assumes each unique cell type secretes the same concentration of cytokines per timestep for all relevant cytokines to drive model agent decisions. Each computational timestep represents 15 minutes of real-world time. We assume that this is of sufficient resolution to accurately reproduce immune cell agent behaviors during regeneration.

**Table 1.**
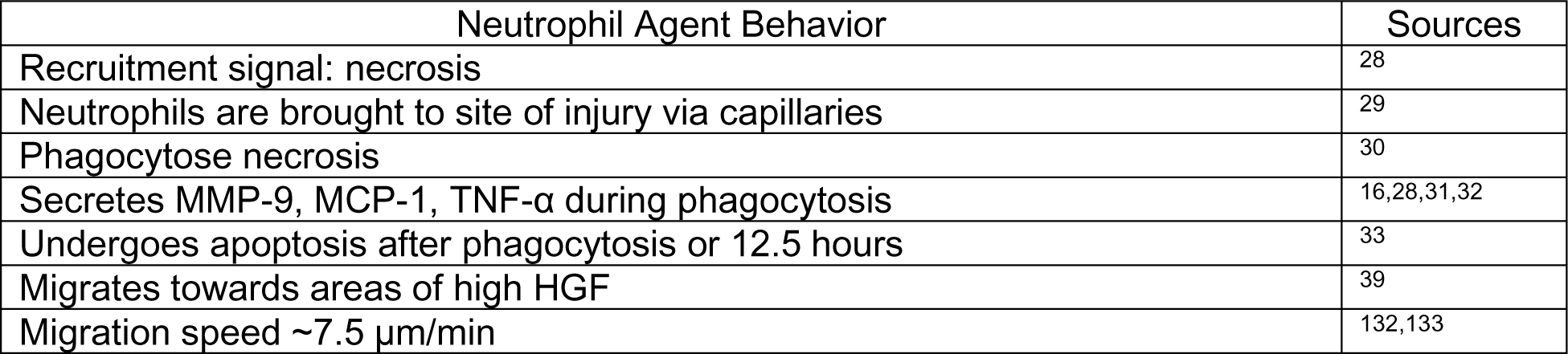
Neutrophil Agent Rules.

**Table 2.** Macrophage Agent Rules.

| Macrophage Agent Behavior | Sources |
| --- | --- |
| Initial count: 1 resident macrophages per 5 myofibers | 34 |
| Recruitment signal: MCP-1 | 38,42 |
| Monocytes are brought to the site of injury via microvessels | 41 |
| Resident macrophages secrete MMP-9, MCP-1, and TNF-α and chemotax along MCP-1 and HGF | 35–40 |
| Monocytes chemotax along MCP-1, VEGF-A, and TGF-β | 42–44,134 |
| Monocyte migration speed $\sim 4 \mu\text{m}/\text{min}$ | 135 |
| M1 macrophages secrete VEGF-A, MMP-9, and TNF-α and chemotax along MCP-1 | 136–139 |
| Monocytes, resident and M1 macrophages phagocytose apoptotic neutrophils and necrosis | 45–47 |
| Monocytes and macrophages secrete MMP-9, HGF, TGF-β, and IL-10 during phagocytosis | 16,48–52 |
| Monocyte transitions to M1 occurs when TNF-α threshold is met or based on literature means and standard deviation properties | 51,53 |
| M1 transition into M2 is mediated by the amount of IL-10 and the amount the M1 has phagocytosed | 16,51,54,55 |
| M2 macrophages secrete TGF-β and IL-10 and chemotax along MCP-1 | 16,38,56,140 |
| Macrophages can proliferate following the transition to the anti-inflammatory (M2) state | 51 |
| Macrophage migration speed $\sim 0.62 \mu\text{m}/\text{min}$ | 135 |
| Macrophages apoptose in a Poisson distribution | 141 |

**Table 3.** SSC Agent Rules.

| SSC Agent Behavior | Sources |
| --- | --- |
| Initial count: 1 SSC per 4 fibers | 15,57 |
| Recruitment signal: HGF + MMP-9 - TGF-β | 15,58–61 |
| Activation signal: HGF | 15,61–64 |
| Activated SSCs secrete MCP-1 and VEGF-A | 42 |
| Activated SSCs migrate towards areas of high MMP-9 | 59,142 |
| Myoblasts migrate towards high TNF-α | 143 |
| Division signal: TNF-α + VEGF-A - TGF-β | 15,60,144,145 |
| Differentiation signal: 3*IL-10 - HGF – TNF-α - TGF-β | 15,54,146–148 |
| Activated SSCs differentiate to myoblasts, myoblasts to myocytes, and myocytes to myotubes/myofibers | 65–67 |
| Differentiated myocytes fuse at damaged fiber edge or fuse together to form new, immature myotubes | 27,68–70 |
| 50% cell divisions are symmetric, 50% asymmetric | 15,149,150 |
| Division probability decreases with each cell division; 1st division 85%; 2nd 65%; 3rd 20% | 15,151 |
| VEGF-A and macrophages nearby can block apoptosis | 42,73,74 |
| TGF-β triggers apoptosis | 75 |
| Time to divide: 10 hrs | 15,151,152 |
| Migration speed $\sim 0.94 \mu\text{m}/\text{min}$ | 153 |
| Return activated SSCs to quiescence without sustained HGF | 61 |

**Table 4.** Fibroblast Agent Rules.

| Fibroblast Agent Behavior | Sources |
| --- | --- |
| Initial count: 1 fibroblast per every 2 fibers | 15,76 |
| Activation signal: TGF-β | 154 |
| Fibroblasts move to low collagen ECM | 15,79 |
| Fibroblasts secrete TNF-α, TGF-β, MMP-9, VEGF-A. Collagen is secreted at low density ECM | 15,80–85 |
| Fibroblast division signaled by SSC division | 15,76 |
| Division probability decreases with each cell division; 1st division 100%; 2nd 25%; 3rd 6% | 155 |
| Fibroblast differentiation to myofibroblasts with extended TGF-β exposure | 15,86,87 |
| Myofibroblasts secrete double the amount of collagen and secretion is not dependent on collagen density | 15,88 |
| Fibroblasts apoptose with sustained exposure to TNF-α | 15,89 |
| Fibroblast migration speed $\sim 0.73 \mu\text{m}/\text{min}$ | 156 |
| Sufficient TGF-β can block fibroblast apoptosis | 15,89 |

**Table 5.**
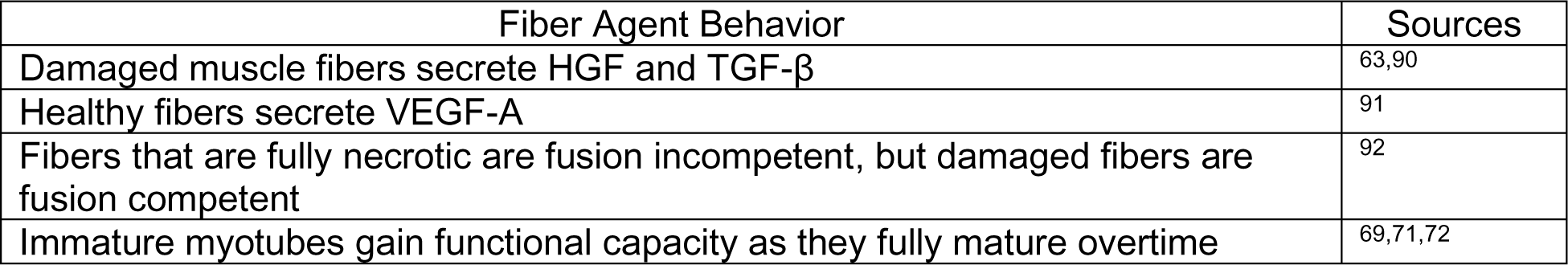
Fiber Agent Rules.

**Table 6.**
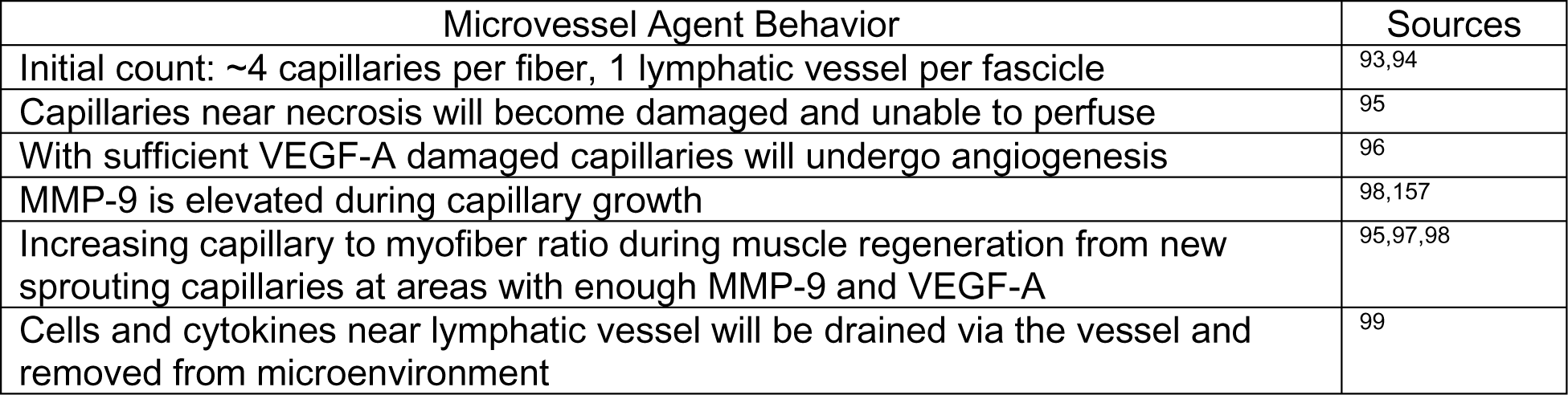
Microvasculature Rules.

| Microvessel Agent Behavior | Sources |
| --- | --- |
| Initial count: $\sim 4$ capillaries per fiber, 1 lymphatic vessel per fascicle | 93,94 |
| Capillaries near necrosis will become damaged and unable to perfuse | 95 |
| With sufficient VEGF-A damaged capillaries will undergo angiogenesis | 96 |
| MMP-9 is elevated during capillary growth | 98,157 |
| Increasing capillary to myofiber ratio during muscle regeneration from new sprouting capillaries at areas with enough MMP-9 and VEGF-A | 95,97,98 |
| Cells and cytokines near lymphatic vessel will be drained via the vessel and removed from microenvironment | 99 |

**Table 7.** Model parameters of spatial mechanisms.

| Parameter | Value | Source/Justification |
| --- | --- | --- |
| *Volume parameters* |  |  |
| Target volume ($V_{target}$) neutrophil | 12 | Chosen for an average cell diameter of $12 \mu\text{m}$<sup>158</sup> |
| Target volume ($V_{target}$) SSC | 10 | Chosen for an average cell diameter of $10 \mu\text{m}$<sup>159</sup> |
| Target volume ($V_{target}$) macrophage | 21 | Chosen for an average cell diameter of $21 \mu\text{m}$<sup>160</sup> |
| Target volume ($V_{target}$) monocyte | 8.5 | Chosen for an average cell diameter of $8.5 \mu\text{m}$<sup>161</sup> |
| Target volume ($V_{target}$) fibroblast | 15 | Chosen for an average cell diameter of $15 \mu\text{m}$<sup>162</sup> |
| Volume multiplier ($\lambda_{volume}$) | 50 | Volume constraint to maintain target<sup>23</sup> |
| *Diffusion coefficients* |  |  |
| HGF | 66.38 $\mu\text{m}^2/\text{s}$ | Estimated diffusivity within the ECM accounting for baseline GAGs and collagen<sup>101</sup> |
| MMP-9 | 63.40 $\mu\text{m}^2/\text{s}$ | |
| MCP-1 | 189.27 $\mu\text{m}^2/\text{s}$ | |
| VEGF-A | 112.10 $\mu\text{m}^2/\text{s}$ | |
| TGF-β | 90.33 $\mu\text{m}^2/\text{s}$ | |
| TNF-α | 138.95 $\mu\text{m}^2/\text{s}$ | |
| IL-10 | 135.17 $\mu\text{m}^2/\text{s}$ | |
| *Chemotaxis parameters $\lambda_c$* |  |  |
| Neutrophils | 750 | Chosen for a cell velocity between 1-20 $\mu\text{m}/\text{min}$<sup>132</sup> |
| Macrophage | 9.3 | Chosen for a cell velocity around 0.62 $\mu\text{m}/\text{min}$<sup>135</sup> |
| Monocyte | 75 | Chosen for a cell velocity around 4 $\mu\text{m}/\text{min}$<sup>135</sup> |
| SSC | 11.3 | Chosen for a cell velocity around 0.94 $\mu\text{m}/\text{min}$<sup>153</sup> |
| Fibroblast | 23 | Chosen for a cell velocity around 0.73 $\mu\text{m}/\text{min}$<sup>8</sup> |

### Neutrophil Agents

Neutrophils are recruited through capillaries to sites of necrotic tissue (Table 1). Neutrophils move to areas of necrotic tissue with high concentrations of HGF by chemotaxing along the HGF gradient to reach areas of necrosis^28,29^. Neutrophils phagocytose necrotic tissue and facilitate remodeling into ECM with low-collagen density. During phagocytosis, neutrophils secrete MMP-9, MCP-1, and TNF-α^30,28,31,32^. Individual neutrophil agents apoptose after phagocytosing two necrotic cells (based on calibration) or 12.5 hours after their recruitment^33^.

### Macrophage Agents

Resident macrophages are distributed randomly throughout the tissue at a ratio of 1 macrophage per 5 myofibers at model initialization and secrete MCP-1^34^ (Table 2). Resident macrophages chemotax along MCP-1 and HGF chemical gradients and secrete MMP-9, TNF-α, and MCP-1 during simulation^35–40^. After tissue injury, monocytes are recruited through healthy capillary microvasculature and chemotax along MCP-1, VEGF-A, TGF-β^41,42–44^. Monocytes infiltrate into the tissue if the MCP-1 concentration is above a specified threshold at a capillary site. Resident macrophages, monocytes, and the M1 macrophages differentiated from monocytes may phagocytose areas of necrotic tissue and apoptotic neutrophil agents^45–47^. During phagocytosis, these agents secrete MMP-9, HGF, TGF-β, and IL-10^48–52^.

Monocytes transition to M1 polarized macrophages when the monocyte agent experiences a large enough TNF-α concentration or if enough time has passed that a predefined transition time threshold is met. Each monocyte agent at creation has a defined transition time sampled from a gaussian distribution with mean and standard deviation (SD) set to reproduce literature-defined populations of M1 macrophages over time^51,53^.

M1 macrophages may transition to M2 macrophages if the M1 macrophage agent experiences an IL-10 concentration that exceeds a threshold value or if the M1 macrophage has phagocytosed enough to meet a calibrated threshold value (as discussed in *Model Calibration*)^51,54,55^. Following the transition to the anti-inflammatory phenotype, the M2 macrophages can proliferate, secrete TGF-β and IL-10, and chemotax along an MCP-1 gradient^38,51,56^.

### SSC Agents

The model is initialized with 1 quiescent SSC per every 4 fibers and upon injury^57^. Additional SSCs are recruited based on the amount of HGF, MMP-9, and TGF-β^58–61^ (Table 3). For SSC activation there has to be enough HGF at the location of the quiescent SSC to induce activation^61–64^. The SSCs also chemotax up the MMP-9 gradient, removing some of the MMP-9 as they move along it. Activated SSCs can also undergo symmetric or asymmetric division and differentiation given that the required cytokine signaling is met locally. Activated SSCs differentiated into myoblasts and myoblasts differentiate into myocytes^65–67^. Myocytes can fuse to other myocytes to form new myotubes or fuse to fibers as long as the fiber is not fusion incompetent (i.e., fully necrotic)^27,68–70^. Maturation of myotubes is required for fusion of additional myocytes to the new fiber^69,71,72^. If the damage signal is not sustained, activated SSCs return to quiescence. If there is enough TGF-β to induce apoptosis and not enough VEGF-A or macrophages nearby to block it, the SSC undergoes cell death and leaves the simulation^42,73–75^.

### Fibroblast Agents

For model initialization, fibroblasts are randomly placed within the ECM at a population size that is proportional to the number of fibers^76^ (Table 4). Fibroblasts are activated based on the concentration of TGF-β around the fibroblast^77,78^. Fibroblasts include an additional expression in their effective energy function that directs their migration towards areas of low-density collagen ECM ^79^. Specifically, fibroblasts can form spring-like links to drag them towards areas of low-density ECM which are implemented with the relation *λ_ij_* (*l_ij_* − *L_ij_*)^2^ where *λ_ij_* denotes a hookean spring constant of a link between cells *i* and *j, l* represents the current distance between the centers of mass between the two cells (in our case, fibroblast and low collagen ECM), and *L* is the target length of the spring-like link. In addition to the cytokines secreted by fibroblasts (Table 4), collagen is secreted at low-density collagen ECM^80–85^. Fibroblasts divide when they are near dividing SSCs and can differentiate into myofibroblasts with extended exposure to TGF-β^76,86,87^. The myofibroblasts can secrete more collagen regardless of the ECM density^88^. Fibroblasts can undergo apoptosis if there are adequate levels of TNF-α at the site of the cell but it can be blocked if there is sufficient TGF-β^89^.

### ECM Agents

ECM elements surround the fiber elements and are assigned a collagen density parameter which varies based on the amount of necrotic tissue removed and the extent of fibroblast/myofibroblast collagen secretion. When necrotic elements are removed, the phagocytosing inflammatory cells secrete MMP-9s which degrade some of the collagen within that section of the ECM, thereby causing that element to have a lower collagen density^28^. The collagen density of the ECM alters the diffusivity of the secreted factors, and fiber placement is dependent on the collagen density (discussed below). The fibroblasts help rebuild the ECM by secreting collagen on low collagen density ECM elements^80^. Myofibroblasts can secrete collagen on any ECM element and if prolonged results in high density collagen elements, representing a fibrotic state.

### Fiber and Necrotic Agents

Upon model initialization, a portion of the muscle fiber agents are converted to necrotic fibers based on the user prescribed injury. Fibers that reach a damaged threshold became fully necrotic whereas those surrounding the area of necrosis were damaged but not fully apoptotic cells. Healthy fiber elements secrete VEGF-A, and necrotic elements secrete HGF and TGF-β^63,90,91^ (Table 5). Phagocytosing agents chemotax along those gradients to clear the necrosis, but before a new fiber can be deposited, the collagen has to be restored so that there is a scaffold to hold the fiber in place^34^. Fully necrotic fibers are fusion incompetent and require myocyte-to-myocyte fusion to form a new myofiber and require maturation before additional myocyte fusion^69,71,72^. Damaged fibers are regenerated by myocytes fusion to the healthy fiber edge^92^.

### Capillary and Lymphatic Agents

The muscle fascicle environment includes approximately 4 capillaries per fiber and 1 lymphatic vessel^93,94^ (Table 6). The model defines perfused capillaries as capillary agents that can transport neutrophils and monocytes into the system proportional to the concentration of recruiting cytokines^29,41^. The neutrophils and monocytes are added to the simulation at the lattice sites above capillaries (within the cell layer Fig. 2B) and chemotax along their respective gradients. The recruitment of the neutrophils and monocytes are distributed among the healthy capillaries with a higher affinity for capillaries at locations with higher concentrations of HGF and MCP-1, respectively. Under physiologically reasonable chemotactic gradient conditions, the recruited immune cells dispersed efficiently, with no aggregation. Capillaries that are neighboring areas of necrosis become non-perfused and therefore are unable to transport cells into the microenvironment until regenerated^95^. Angiogenesis can occur as long as there is enough VEGF-A present at the non-perfused capillary^96^. Similar to published studies, there is an increase in the capillary-to-myofiber ratio during muscle regeneration, which is due to the formation of new capillary sprouts modulated in part by local MMP-9 and VEGF-A levels^95,97,98^.

The lymphatic vessel uptakes cytokines at lattice locations corresponding to the lymphatic vessel and will remove cells located in lattice sites neighboring those corresponding to the lymphatic vessel^99^. In addition, we have included a rule in our ABM to encourage cells to migrate towards the lymphatic vessel utilizing CompuCell3D External Potential Plugin^100^. The influence of this rule is inversely proportional to the distance of the cells to the lymphatic vessel.

### Binding, Diffusivity, and Collagen Density

For many of the agent behaviors described above, there are associated binding events that play key roles in regulation of the cytokine fields. Any cytokine dependent behavior is coupled with removal of a portion of that cytokine once the behavior is initiated. For example, upon SSC activation the amount of HGF required to activate is taken up by the SSC and removed from the cytokine field to simulate the ligand binding and endocytosis resulting from SSC activation. Similar binding events were modeled for SSC and fibroblast division and differentiation, macrophage transitions, cell apoptosis, and chemotaxis along a cytokine gradient.

Due to limited data availability quantifying the diffusion constants of the modeled cytokines in the context of the tissue microenvironment (which includes diffusion-altering elements including collagen and glycosaminoglycans (GAGs)), we applied a diffusivity estimation technique^101^. To do so, previously developed methods (equation 1) were applied to account for the combined effects of collagen and GAGs (Table 7)^101^. The expression includes the radius of the cytokine (r_s_), the radius of the fiber (r_f_), the volume fraction (*ϕ*), *D* and *D*_∞_ are the diffusivities of the cytokines in the polymer solution and in free solution, respectively. This estimation technique allowed for consistent conditions for cytokine diffusion calculations and fluctuations based on changes in collagen density within the model.

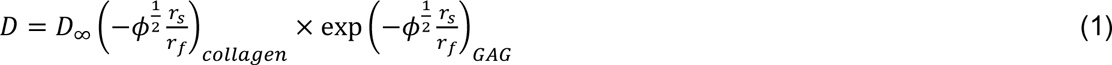

Throughout the model simulation, the diffusivity is recalculated with the updated collagen volume fraction, as the collagen density changes throughout the microenvironment. This allows the changes in collagen density within the ECM to be reflected in the diffusion rate of each of the cytokines in the model.

### Model Calibration

Known parameters were fixed to literature values, and uncertain parameters were calibrated by comparing simulation outcomes to published experimental data. Calibration data included published findings from injury models that have synchronous regeneration after tissue necrosis (i.e., cardiotoxin, notexin, and barium chloride)^97^. The metrics that were used to calibrate the model included time-varying CSA^102^, SSC counts^76^, and fibroblast counts^76^. These metrics were used for calibration because of their key roles in the regeneration of muscle and the complex interplay between these outputs. Cell count data were normalized by the number of cells on the day of the experimental peak to allow for comparison between experiments and simulations. For CSA, the experimental and model outcomes were normalized using fold-change from pre-injury to compare model-simulated with experimental CSA, as percent change from baseline is commonly used experimentally^103,104^. Model cell counts were normalized by the number of cells at the peak time point in the experimental data. SSC and fibroblast counts were normalized to day 5. Neutrophil counts were normalized to day 1. Total macrophage, M1, and M2 counts were normalized to day 3. The capillaries were normalized to fiber area, as done in the experimental data.

Initial ranges for the 52 unknown parameters were determined by literature review or by running the model to test possible upper and lower thresholds for parameters (Supplemental Table 1). To narrow the parameter ranges beyond those initial ranges, we used a recently published calibration protocol, CaliPro, which utilizes parameter density estimation to refine parameter space and calibrate to temporal biological datasets^20^. CaliPro was selected as the calibration method because it is model-agnostic which allows it to handle the complexities of stochastic models such as ABMs, selects viable parameter ranges in the setting of a very high-dimensional parameter space, and circumvents the need for a cost function, a challenge when there are many objectives, as in our case. Briefly, Latin hypercube sampling (LHS) was used to generate 600 samples which were run in triplicate. These runs were then evaluated against a set of pass criteria, and the density functions of the passing runs and failing runs were calculated (Supplemental Table 2). Parameter ranges were narrowed by alternative density subtraction (ADS), where the new ranges were determined by the smallest and largest parameter values where the density of passing is higher than the density of failing. The sensitivity of the model outputs to the parameters was examined using LHS in combination with partial rank correlation coefficient (PRCC)^105^. LHS/PRCC methods have been used for various differential equation models and ABMs^106^. PRCC was computed using MATLAB to determine the correlation between ABM parameters (i.e. cytokine threshold for activation) and the ABM output (i.e. fibroblast cell count). Correlations with a p-value less than 0.05 were assumed to be statistically significant. This helped refine initial parameter bounds as well as make model adjustments based on the parameter dynamics elucidated from PRCC. This process of sampling parameter ranges, evaluating the model, and narrowing parameter ranges was repeated in an iterative fashion while updating pass criteria until a parameter set was identified that consistently met the strictest criteria (Supplemental Fig. 1). The final passing criteria were set to be within 1 SD of the experimental data for CSA recovery and 2.5 SD for SSC and fibroblast count. These criteria were selected so that the model followed experimental trends and accounted for both model stochasticity and experimental variability in datasets that had narrower SDs for certain timepoints. Early iterations had a wide parameter range to avoid missing portions of the realistic parameter space. At first, narrowing the parameter space increased passing simulations, but upon reaching the ideal parameter space, further narrowing eliminated viable parameters, resulting in fewer passing runs. Following 8 iterations of narrowing the parameter space with CaliPro, we reached a set of parameters that had fewer passing runs than the previous iteration. We then returned to the runs from the prior iteration and set the bounds such that all 3 runs from the parameter set fell within the final passing criteria. The final parameter set was run 100 times to verify that the variation from the stochastic nature of the rules did not cause output that was inconsistent with experimental trends.

### Model Validation

We compared model outputs (M1, M2, and total macrophage counts^97,107^, neutrophil counts^108^, and capillary counts^102^) that were kept separate from the calibration criteria with published experimental data to verify that these outputs followed trends from the experimental data without requiring extra model tuning. In addition, we also altered various model input conditions (cell input conditions, cytokine dynamics, and microvessel dynamics) to simulate an array of model perturbations (Table 8) which allowed comparison of a set of model outputs with separate published experiments. For example, we simulated an IL-10 knockout condition by eliminating IL-10 secretion and adjusting the diffusion and decay parameters so that the concentration of IL-10 throughout the simulation was reduced, decreasing the behaviors driven by the cytokine as a result of the KO condition. One hundred replicates of each model perturbation were performed, and perturbation outputs were compared with control simulation outputs via a two-sample t-test with a significance level of 0.05. We were then able to compare how the model outputs aligned with published experimental findings to determine if the model could capture the altered regeneration dynamics.

**Table 8.** Model perturbation input conditions and corresponding published experimental results.

| **Perturbation** | **Specific model conditions** | **Published outcomes** |
| --- | --- | --- |
| IL-10 knockout | Adjust diffusion and decay parameters so IL-10 is removed from the system | Attenuates shift to M2, disrupted SSC differentiation, slowed regeneration<sup>163</sup> |
| Neutrophil depletion | Lower neutrophil recruitment proportion | Abundant necrotic tissue 7 days post injury<sup>109</sup> |
| Macrophage depletion | Lower macrophage recruitment proportion | Decreased HGF, increased TGF-β and TNF-α, impaired regeneration<sup>110</sup> |
| MCP-1 Knockout | Adjust diffusion and decay parameters so MCP-1 is removed from the system | Increased necrosis at day 7, lower CSA at day 21, impaired phagocytosis<sup>115</sup> |
| Directed M2 polarization (Anti-inflammatory nanoparticles) | Require less phagocytosis and IL-10 for transition | Improved muscle histology and inflammatory resolution<sup>164</sup> |
| TNF-α Knockout | Adjust diffusion and decay parameters so TNF-α is removed from the system | Impaired recovery at days 5 and 12, increased inflammation<sup>165</sup> |
| Hindered angiogenesis | Increase VEGF-A and MMP-9 threshold required for angiogenesis | Delayed regeneration with toxin injury, and persistent immune cell infiltration with freeze injury<sup>111</sup> |
| VEGF-A injection | Add VEGF-A at specified concentration (100 for low and 1000 relative concentration for high), radius (300 pixels), and timepoint (5 days post injury) | Lower injury area at day 20 post injury with injection 5 days after damage<sup>73</sup> |

### Sensitivity Analysis

A sensitivity analysis was performed using LHS-PRCC to examine the impact of cytokine-related parameters on model outputs of interest. Diffusion coefficients and decay rates for the seven cytokines (HGF, TGF-β, MMP-9, TNF-α, VEGF-A, IL-10, MCP-1) were sampled across a range from 0.1 to 10 times the calibrated value while holding the other parameters constant. Three hundred samples were generated, and these parameter sets were simulated in triplicate. PRCCs were calculated with α=0.05 and a Bonferroni correction for the number of tests every 10 ticks/hours for CSA and cell counts for SSCs, fibroblasts, non-perfused capillaries, myoblasts, myocytes, neutrophils, M1 macrophages, and M2 macrophages.

### In Silico Experiments

To gain insight into the recovery response with altered angiogenesis, we simulated different levels of VEGF-A injections to test how increases in VEGF-A impacted regeneration outcomes. In addition, we simulated conditions of hindered angiogenesis in which damaged capillaries were unable to reperfuse following injury (n = 100 for each simulation condition). Simulations were also conducted to examine correlations between cytokines and their impact on various cell behaviors and regeneration outcomes. Next, a sensitivity analysis was performed to understand how alterations in cytokines influence key metrics of regeneration. LHS-PRCC was used to quantify the impact of cytokine-related parameters (i.e. diffusion rates and decay coefficients) on outputs of interest (CSA, SSC, fibroblasts, non-perfused capillaries, myoblasts, myocytes, neutrophils, M1, and M2). A single time point for each output is summarized in Table 9, and these were chosen at the timepoint when PRCC values were peaking, with complete results available in Supplementary Figure 2.

**Table 9.**
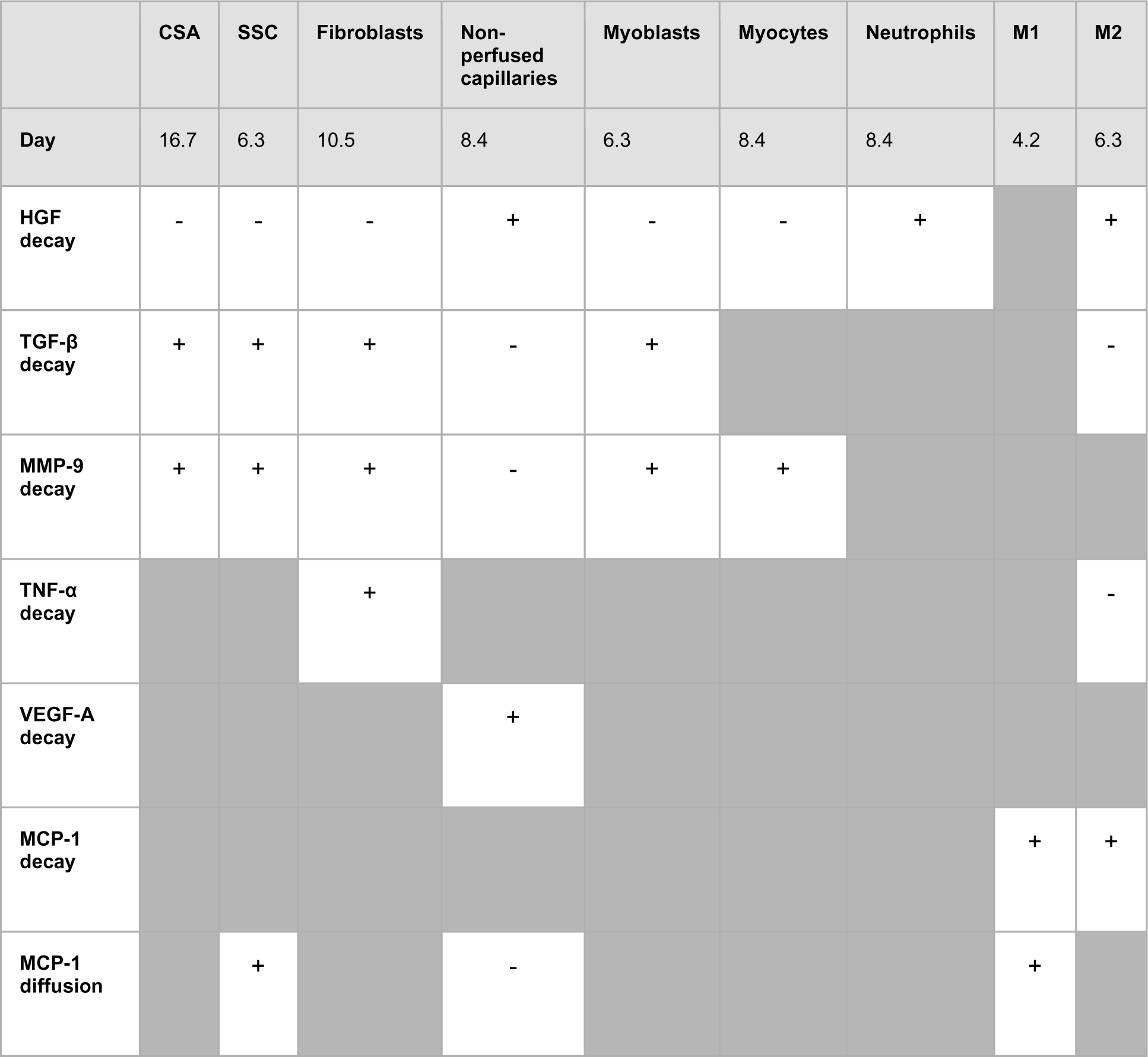
Summary of cytokine sensitivity analysis. Significance was determined with α=0.05, and a Bonferroni correction for the number of tests. + and - represent statistically significant positive and negative correlations, respectively.

|  | CSA | SSC | Fibroblasts | Non-perfused capillaries | Myoblasts | Myocytes | Neutrophils | M1 | M2 |
| --- | --- | --- | --- | --- | --- | --- | --- | --- | --- |
| Day | 16.7 | 6.3 | 10.5 | 8.4 | 6.3 | 8.4 | 8.4 | 4.2 | 6.3 |
| HGF decay | - | - | - | + | - | - | + |  | + |
| TGF-β decay | + | + | + | - | + | | | | - |
| MMP-9 decay | + | + | + | - | + | + |  |  |  |
| TNF-α decay | | | + | | | | | | - |
| VEGF-A decay |  |  |  | + |  |  |  |  |  |
| MCP-1 decay |  |  |  |  |  |  |  | + | + |
| MCP-1 diffusion |  | + |  | - |  |  |  | + |  |

This sensitivity analysis was then used to guide *in silico* experiments based on which cytokine parameters promoted favorable regeneration outcomes (i.e. improved recovery, fewer non-perfused capillaries, increased SSCs). Following individual cytokine parameter alterations, we combined the cytokine alterations based on beneficial outcomes from the initial *in silico* experiments to determine if the benefits would be cumulative.

## Results

### ABM outputs align with calibration and validation data

Following parameter density-based calibration, the unknown parameters were narrowed into a final calibration parameter set (Supplemental table 1). The simulations captured SSC and fibroblast cellular behaviors, as well as CSA outcomes, that aligned with experimental studies (Fig. 3A-C). The model data were consistent with the experimental trends, and the 95% confidence interval was within the SD for all calibration data time points except for SSCs at day 3 (Fig. 3B). Macrophage (total, M1, and M2), neutrophil, and capillary counts, which were not used for model calibration, were found to also be consistent with experimental trends and allowed us to independently validate model outputs (Fig. 3D-H).

**Figure 3.**
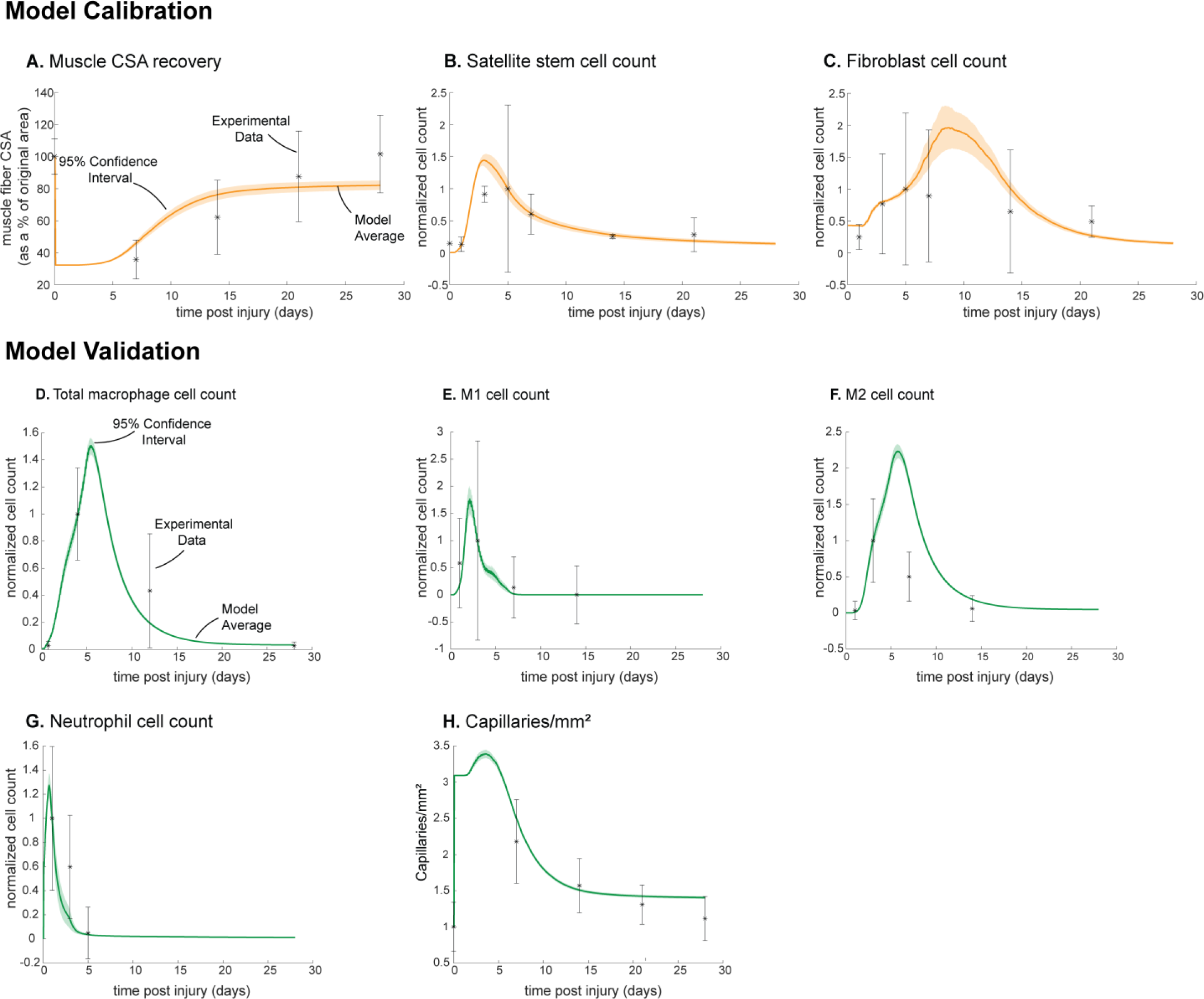
ABM calibration and validation. ABM parameters were calibrated so that model outputs for CSA recovery, SSC, and fibroblast counts were consistent with experimental data (A-C)^76,102^. Separate outputs from those used in calibration were compared to experimental data^97,102,107,108^ to validate the ABM (D-H). Error bars represent experimental standard deviation, and model 95% confidence interval is indicated by the shaded region. Cell count data were normalized by number of cells on the day of the experimental peak to allow for comparison between experiments and simulations.

### ABM perturbations are consistent with published experiments

Overall, the model reproduced findings from multiple studies, replicating how altered conditions lead to both improved and diminished muscle regeneration (Fig. 4). Injections of VEGF-A led to faster CSA recovery, more damaged tissue clearance, and a concentration dependent dose response, consistent with prior studies^73^. Cell depletion simulations predicted decrease in all markers of regeneration, consistent with prior studies^73,109,110^. When simulating hindered angiogenesis conditions, the model aligned with experimental studies showing detriments in CSA recovery, increased neutrophil and macrophage cells, and elevated ECM collagen density, indicating progression of fibrosis within the microenvironment^111^. There were a few cases in which model predictions did not align with published studies. First, simulations of TNF-α KO predicted increased CSA recovery, while experiments measured decreased recovery of CSA. This difference is likely due to the fact that the model did not include cross regulation with interferons which are upregulated with TNF-α KO^112^. Second, macrophage depletion simulations predicted decreased TGF-β concentrations throughout the simulation while experiments measured an initial decrease in concentration followed by increased concentrations at days 7 and 14. This difference may be due to the fact that macrophage depletion was experimentally induced with clodronate-containing liposomes which could have reduced consistency of depletion across the time course and other downstream impacts that were not represented by decreasing macrophages in the model perturbation^110^.

**Figure 4.**
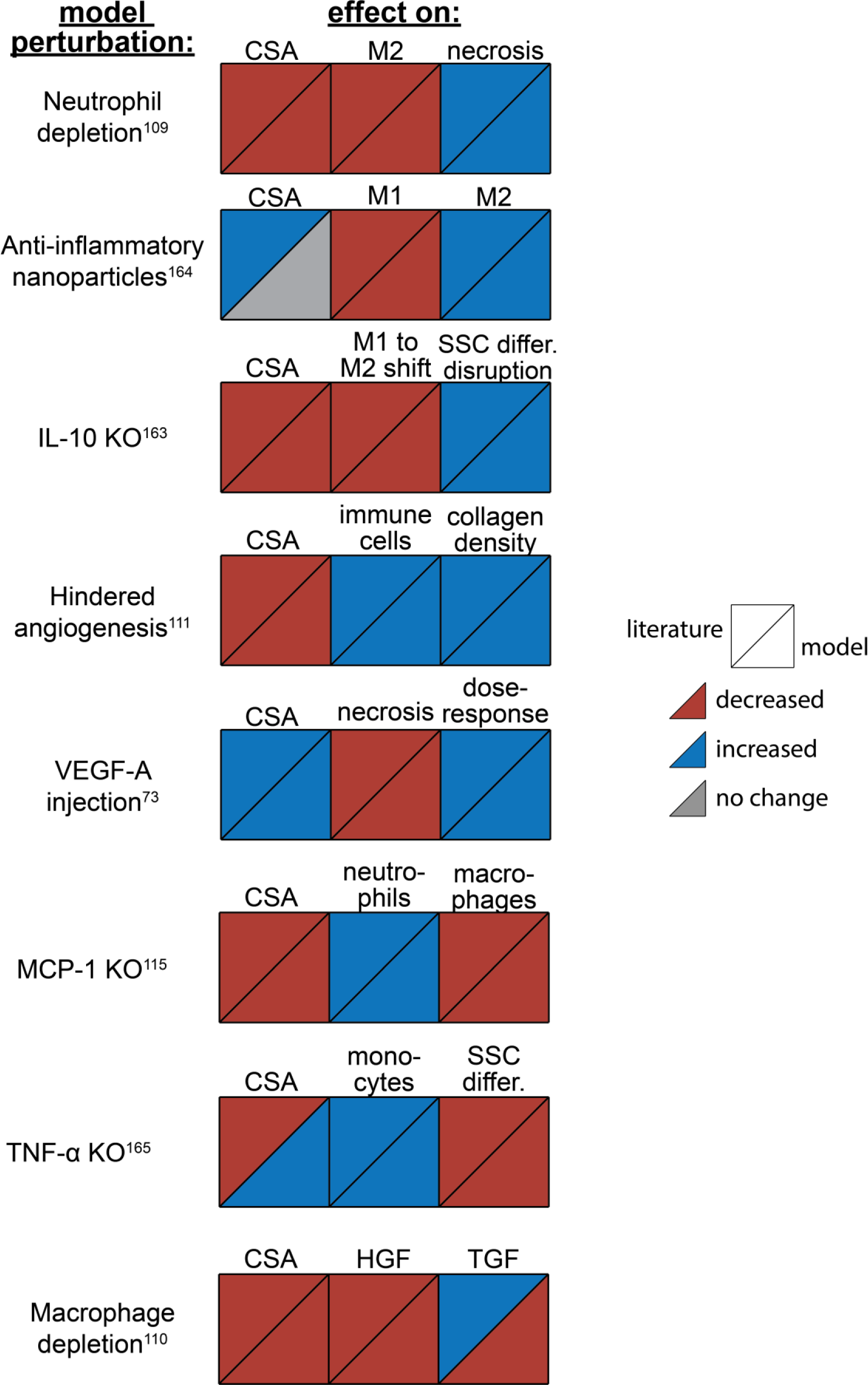
ABM perturbation outputs are compared to various literature experimental results. Each perturbation model output is compared to the available corresponding published result. The top triangles indicate the literature findings and the bottom triangles indicate the model outputs. Red triangles represent a decrease, blue represents an increase, and gray represents no significant change. Time points of comparison were based on which time points were available from published experimental data. Refer to table 8 for model input conditions and supplemental table 6 for information on experimental references.

### Analysis of ABM perturbations leads to new insights regarding cytokine and cell dynamics

The model allowed for new insights into the dynamics of muscle regeneration by providing additional timepoints and metrics to evaluate the response to exogenous delivery of VEGF-A and hindered angiogenesis. VEGF-A levels remained elevated compared to control simulations following the injection at day 5 post injury (Fig. 5A). CSA recovery had the highest increase at 28 days post injury with the high (10^3^) relative concentration delivered) VEGF-A injection followed by the extra high (2×10^3^) relative concentration delivered) injection (Fig. 5B). The medium (750 relative concentration delivered) and low (500 relative concentration delivered) VEGF-A injections had higher CSA recovery 15 days post injury but were not significantly different from the control at day 28. All VEGF-A injections had a higher capillary count and were proportional to the level of VEGF-A injection (Fig. 5C). The impact of VEGF-A injection on peak SSC and fibroblast counts was dependent on dosage amount, with the extra high VEGF-A injection resulting in the largest peaks (Fig. 5E&F). Cytokine concentration trends were similar for all injections, but most peak levels were dosage dependent (Fig. 5G-L). In contrast, HGF levels were elevated from day 5 to day 28 with hindered angiogenesis, as were TGF-β and IL-10 (Fig. 5I & L). MCP-1 concentration had a lower overall peak level with elevated levels from days 21 to 28 (Fig. 5H). Hindered angiogenesis had lower CSA recovery throughout the simulation and did not achieve unaltered regeneration levels (Fig. 5B).

**Figure 5.**
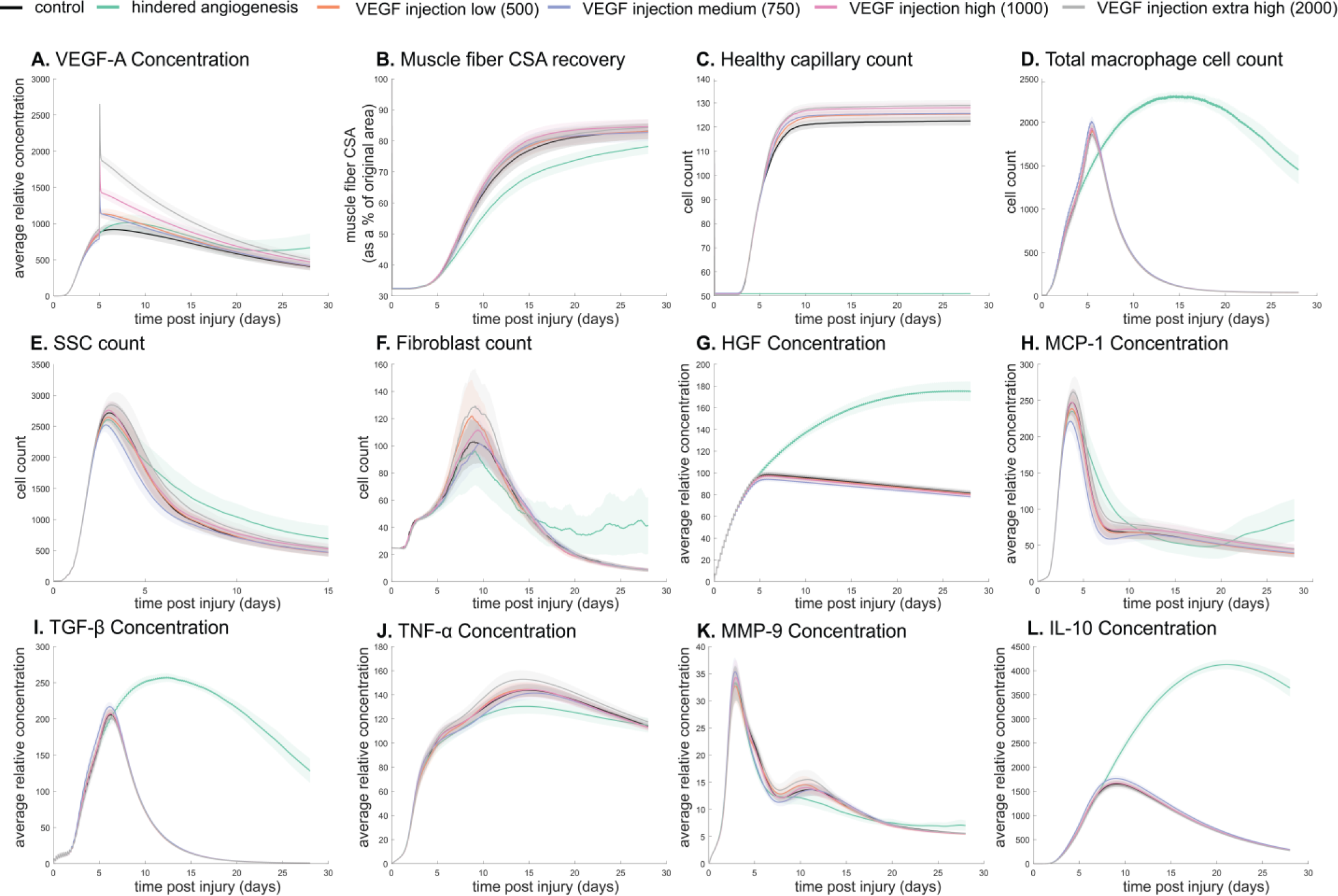
Dose dependent response with VEGF-A injection compared to hindered angiogenesis. VEGF-A concentration response to varied levels of VEGF injection (A). Hindered angiogenesis resulted in slower and overall decreased CSA recovery (B). Capillary count was dependent on VEGF-A injection level (C). Total macrophage count was similar between control and VEGF-A injections perturbations but macrophage count was higher in later time points in the hindered angiogenesis simulation (D). SSC peak varied with VEGF-A injection level and counts were prolonged in the hindered angiogenesis simulations (E). The fibroblast peak was lower for the hindered angiogenesis perturbation and highest with the extra high VEGF-A injection. In contrast to the other simulations, the fibroblast count was trending upwards at later time points in the hindered angiogenesis perturbation (F). HGF levels were consistent between control and VEGF-A injection perturbations but was significantly elevated in the hindered angiogenesis perturbation (G). MCP-1, TGF-β, and IL-10 concentrations were elevated a later stages of regeneration with hindered angiogenesis (H, I, L). TNF-α was elevated with the extra high VEGF-A injection and lower with hindered angiogenesis (J). MMP-9 concentration was lower at the simulation midpoint but elevated at late regeneration stages (K).

Cytokine knockout perturbations revealed crosstalk and temporal interplay between cytokines (Fig. 6). For example, with MCP-1 KO there was an overall increase in cytokine levels for all other cytokines within the microenvironment except for VEGF-A at 12 hours post injury (Fig. 6A). By 7 days post injury TNF-α, TGF-β, IL-10, and MMP-9 had decreased from unaltered regeneration day 7 levels but VEGF-A and HGF were elevated. With TNF-α KO there was a decrease in TGF-β at early timepoints but a strong increase by day 28 (Fig. 6B). Following IL-10 KO there was an increase in TNF-α that peaked at 7 days post injury (Fig. 6C). HGF was slightly decreased throughout and TGF-β was strongly decreased by day 7. MMP-9 was decreased at 12 hours and 28 days post injury but heavily increased at day 7.

**Figure 6.**
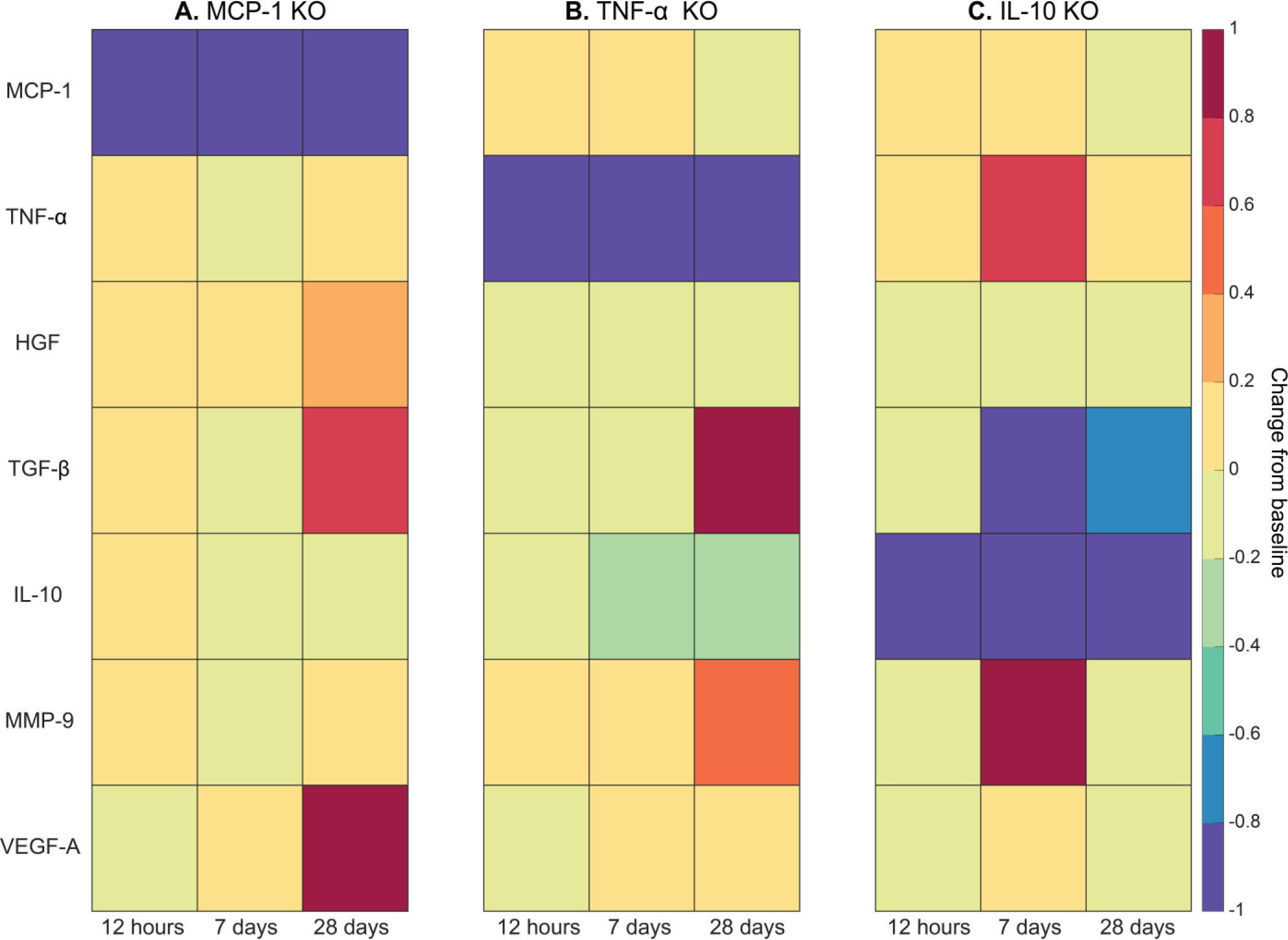
Heatmaps of changes in cytokine concentration at various timepoints throughout regeneration following individual cytokine knockout (KO) demonstrating cross-talk between cytokines. With MCP-1 KO there was an increase in all cytokines except VEGF-A at 12 hours post injury. Over the course of regeneration there was continued increasing elevation of HGF, increases in VEGF-A, and TGF-β decreased at day 7 followed by a strong increase by day 28 post injury (A). In the TNF-α KO simulations, there was an early decrease in TGF-β that shifts to strong increases by day 28. MMP-9 increased throughout the duration, HGF and IL-10 were decreased, VEGF-A lagged in the beginning but was increased during mid to late timepoints (B). Following IL-10 KO there were increases in TNF-α, decreases in HGF and TGF-β, and elevated MMP-9 at day 7 that decreased by day 28 (C).

### Cytokine dynamic analysis leads to new model perturbations that predict improved regeneration

LHS-PRCC of cytokine decay and diffusion parameters elucidated temporal relationships between cytokine parameters and key regeneration metrics, such as positive correlations between CSA and TGF-β and MMP-9 decay (Table 9). Of all cytokine parameters, the model outputs were most sensitive to HGF decay, with all outputs except M1 cell count being significantly impacted. PRCC plots showed that TGF-β and MMP decay were positively correlated and HGF decay was negatively correlated with CSA recovery, with higher significance at timepoints after 12 days (Supplemental Fig. 2). Correlation plots for various cytokine concentrations and regeneration metrics showed trends in cytokine dependent cell behaviors such as the TNF-α concentration that led to heightened fibroblast cell counts as well as the corresponding TNF-α concentration threshold that results in diminished fibroblast response (Supplemental Fig 3). These PRCC trends guided cytokine parameter perturbations to include lower HGF and VEGF-A decay, higher TGF-β, MMP-9, and MCP-1 decay, and higher MCP-1 diffusion because each of the cytokine modifications indicated some form of enhanced regeneration outcome metrics (Supplemental Table 3). All these perturbations except MCP-1 decay show increased CSA, increased healthy capillaries, and increased SSCs (Fig. 7). Finally, a combination of all changes except for MCP-1 decay was simulated. The combined cytokine alteration resulted in the highest CSA recovery (Fig. 7A), as well as increased M1 macrophage counts (Fig. 7B), decreased M2 macrophage counts (Fig. 7C), increased fibroblasts (Fig. 7D) and SSCs cell counts (Fig. 7E). Capillaries regenerated faster in the combined perturbation than under unaltered conditions (Fig. 7F). It is likely that the combination of cytokines perturbed cell dynamics in a manner that promoted regeneration in both the early and later phases. During early regeneration, lower HGF decay, higher TGF decay, and MCP-1 diffusion contributed to increased SSCs while lowered VEGF decay increased angiogenesis. During late regeneration, lower HGF decay and higher MMP decay contributed to an increased anti-inflammatory state and SSC differentiation. The combined cytokine perturbation predicted a 13% improvement in CSA recovery compared to the unaltered regeneration amount at 28 days. The combined cytokine perturbation also had higher peaks in SSC and fibroblast counts than any of the singular cytokine perturbations, indicating the synergistic effects of altering the cytokine dynamics in combination.

**Figure 7.**
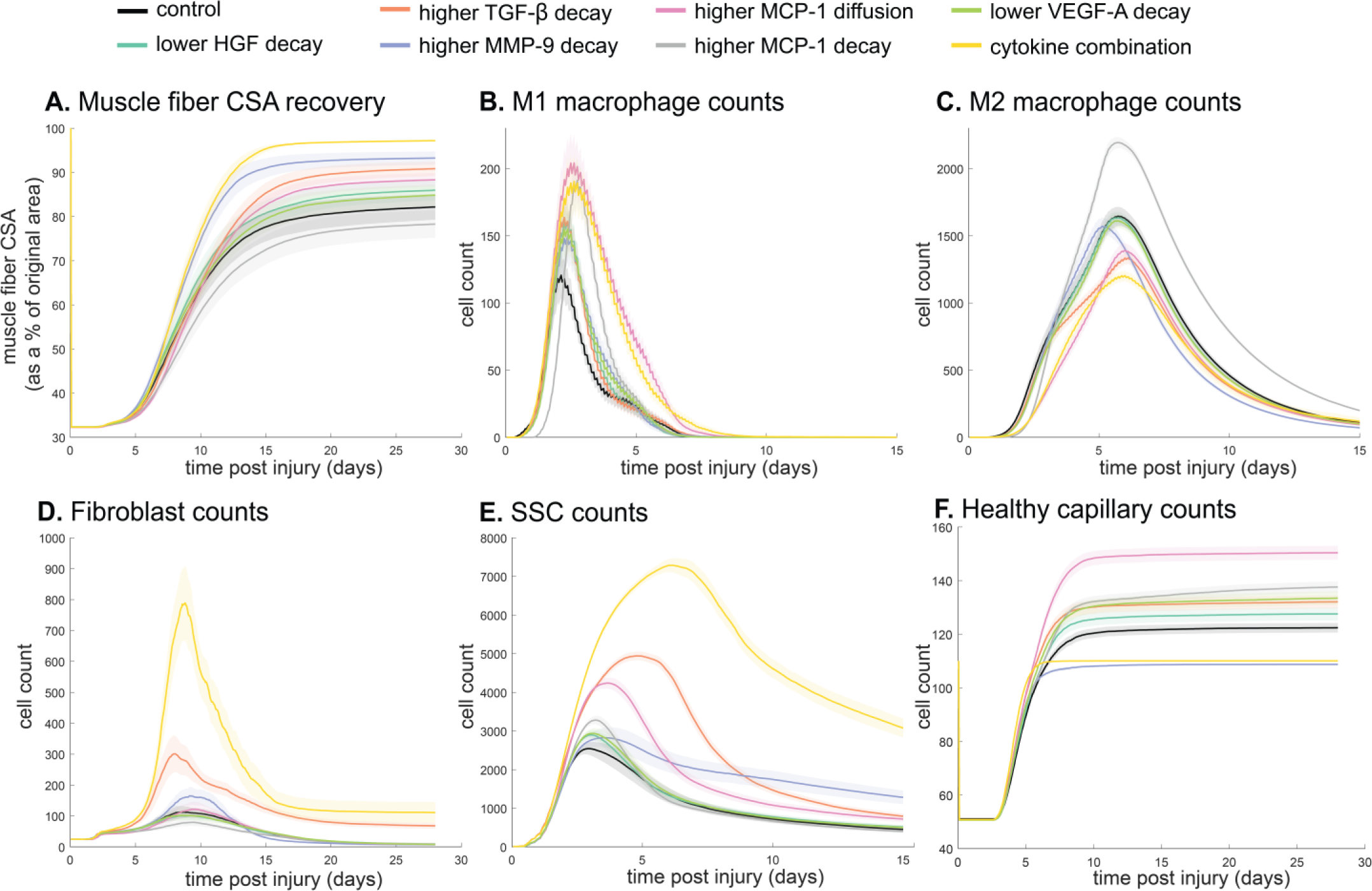
Combined alterations of various cytokine dynamics enhance muscle regeneration outcomes. All tested alterations except higher MCP-1 decay resulted in higher CSA recovery compared to the control (A). M1 cell count was higher for all perturbations with the highest peaks with increased MCP-1 diffusion and the combined cytokine alteration perturbation (B). Higher MCP-1 decay resulted in the largest M2 peak and higher MCP-1 diffusion, higher TGF-β decay, and the combined cytokine alteration had a lower M2 peak than the control (C). Fibroblasts had the largest increase in cell count with the higher TGF-β decay and the cytokine combination perturbations (D). All perturbations resulted in an increased in SSC count with the largest increase resulting from the combined cytokine alteration (E). All perturbations except the combined and higher MMP-9 decay resulted in increased capillaries as a result of additional capillary sprouts (F).

## Discussion

We developed a novel ABM that recapitulates muscle regeneration and, unique from prior work, includes spatial interactions between cytokines and the microvasculature based on relevant literature^8,15,16^. The creation of the model provides a more controlled environment for studying muscle regeneration, reducing error and variation commonly encountered with *in vivo* experiments. Model predictions aligned with experimental data under various altered inputs. Through *in silico* experiments, we gained new insight into how the combination of key cytokine dynamic alterations could increase SSC cells and enhance CSA recovery. The ability for altered cytokine concentrations to change regeneration outcomes is consistent with studies that have found enhanced muscle recovery with delivery of platelet-rich plasma (PRP) which contain VEGF and TGF-β^113^. These model perturbations allow development of hypotheses and can provide the basis for future experiments and potential therapeutic interventions such as plasminogen activators to alter cytokines dynamics to enhance muscle recovery.

### ABM provides biological insight on nonlinear effects of cytokine levels

The ABM offers valuable insights into the muscle regeneration dynamics under various altered conditions, elucidating the complex interplay of cytokines, angiogenesis, and cell behaviors. Systematic simulations reveal critical thresholds, nonlinear effects, and synergistic cytokine combinations impacting regeneration. Perturbations varying VEGF-A injection doses showed increased CSA recovery up to a threshold (high VEGF-A injection simulation), beyond which further improvements in CSA recovery cease. Cytokine KO simulations revealed the complex nature of the relationship between cytokines; removal of one cytokine from the system has a cascading temporal impact. Relationships between cytokines and cellular outputs exhibit nonlinear effects, as seen with the limited impact of elevated HGF on CSA recovery beyond a threshold and the non-monotonic relationship between TNF-α and fibroblast counts (Supplemental Fig. 3). Further analysis revealed that specific combinations of cytokine perturbations could enhance regeneration beyond singular cytokine interventions. For example, a combined intervention of: 1) decreasing HGF and VEGF-A decay, 2) increasing TGF-β and MMP-9 decay, and 3) increasing MCP-1 diffusion enhanced muscle regeneration. Prior studies have shown that individually, increased HGF^114^, VEGF-A^73^, and MCP-1^115^ stimulate muscle regeneration whereas reduced TGF-β^116^ and MMP-9^117^ stimulate muscle regeneration. The model suggests that combined alterations have a stronger regenerative effect than individual cytokine changes, enhancing muscle recovery through distinct mechanisms—increasing healthy capillaries, SSC counts, and reducing inflammatory cells.

Cytokine modifications intended to enhance muscle recovery can have clinical relevance and have been studied in various settings. For example, synthetic biomaterials coated with IL-4 have been implanted as a cytokine delivery vehicle and were successful in increasing M2 cells within the muscle^118^. Cytokine antagonist have been successful at promoting muscle regeneration, seen in prior work with anti-IL-6^119^. Studies have also shown that activation of plasmin is able to induce the release of ECM bound VEGF, increasing angiogenesis^14,120^. Due to the complex network of cytokines, studies that deliver simple modulation of one or two cytokines typically have an insufficient response to generate appreciable improvements. This suggests that using a combination of biologic and synthetic biomaterials to modulate multiple cytokines is necessary, which aligns with our findings^118^. Multiple cytokines have been modulated through the use of PRP which contains VEGF-A and an array of other cytokines, but PRP has had mixed success in a clinical setting^121^. Our model has the capability to test and optimize various combinations of cytokines, along with exploring different temporal schedules for delivering specific treatments. For instance, it can predict whether modified combinations of cytokines prove beneficial at specific timepoints, aiding in the development of optimal treatment compositions aligned with the temporal dynamics of the regeneration cascade. These predictions provide novel concepts for future experiments and potential interventions. For example, the predictions from the model suggest that interventions that combine activation of plasmins for bound VEGF release^14,120^ with delivery of synthetic biomaterials coated with HGF^122^, TGF-β antagonist^123^, nuclear factor-kappa B inhibitory peptide to inhibit MMP-9^124^, and recombinant MCP-1 hydrogels^125^ to alter diffusion rate would result in improved regeneration outcomes.

### Advancements from prior muscle regeneration models

Previous studies have employed computational models to investigate muscle regeneration across diverse contexts, such as Duchenne muscular dystrophy and volumetric muscle loss^8,15^. Earlier muscle regeneration ABMs from our group have been used to test the effects of priming muscle with inflammatory cells prior to injury^16^. While these models laid the foundation for simulating muscle adaptations, they were constrained by limited diffusion capabilities and an absence of critical features related to microvessel growth and remodeling throughout the regeneration process. Similarly, other ABMs from our group have examined altered microenvironments, but their omission of spatial cytokine diffusion hindered comprehensive representation of cell behaviors pivotal to regeneration^8,15^. Recently, new ABMs have been published that focus on cerebral palsy and the impact of injury type on eccentric contraction-induced damage^17,18^.

The model presented here provides advancements over prior models in three areas: 1) explicit modeling of cytokine-specific diffusion and decay that depends on the ECM environment, 2) addition of microvasculature, and 3) incorporation of a robust and rigorous calibration and validation process. The addition of microvessel growth and remodeling dynamics empowers investigations into how interventions impact angiogenesis during regeneration, thereby influencing muscle recovery outcomes. By considering the intricate relationship between microvessels and regeneration, our model opens avenues for evaluating the effects of interventions on the broader recovery process. Secondly, understanding how cytokines influence cell behaviors at different times during regeneration is crucial for determining optimal treatment targets and dosing. While cytokine dynamics can be altered experimentally, doing so is expensive and time consuming^12,14^ so exploring many combinations of alterations would be practically infeasible. Our model incorporates decay and diffusion dynamics of a subset of cytokines to allow testing of far more alterations in cytokines than would be reasonable to conduct experimentally. Lastly, we leveraged the CaliPro technique for parameter density estimation-based calibration and LHS-PRCC to gain biological insight by analyzing how altered microenvironmental parameters could benefit regeneration outcomes. This approach of implementing parameter identification to guide model perturbations demonstrates the capabilities of the model as a novel tool for generating new hypotheses and identifying mechanisms to target for enhanced regeneration outcomes.

Our model predictions are generally consistent with these prior models, with added biological complexity that has yielded several new important insights. For example, simulation of hindered angiogenesis predicted a decrease in SSCs leading to poor CSA recovery, similar to how lower SSC counts resulted in lower CSA recovery in perturbations in both healthy and DMD simulations^15^. Our model provides additional understanding about the corresponding spatial cytokine changes that ultimately result in modulation of SSC dynamics within the microenvironment. The additional model advancements incorporated address prior muscle regeneration modeling gaps in understanding of how angiogenesis alters recovery outcomes as well as the response of complex spatial cell and cytokine dynamics.

### Limitations and future work

There are some important limitations of this study that should be discussed. First, the model does not include all cell types and cytokines that are known to influence muscle regeneration and does not account for cytokine subtype or differences between endogenous and exogenous cytokines. These cells and cytokines likely have redundant functions, given the model effectively captures muscle regeneration using the included cells and cytokines. Second, the model does not currently represent hypertrophy during regeneration, which restricts CSA recovery from surpassing 100%; however, the cell dynamics it portrays remain consistent with those observed in studies that lead to hypertrophy following injury. Third, we assume a 2D cross-section based on similar ABMs that have explored the relations of 2D to 3D simulations. These studies found that the diffusion accuracy is not greatly varied and that 2D is sufficient to predict the same mechanisms seen in 3D simulations^126,127^. To determine the robustness of the 2D initial cross-section, preliminary testing has shown that the initial spatial configuration can be altered and still achieve similar results (Supplemental Fig. 5), but further examination is needed to determine sensitivity to numerous configurations. Fourth, the calibration and validation dataset integrated multiple datasets from diverse sources. We acknowledge inherent limitations arising from variations in sample sizes and experimental techniques across sources. Fifth, it is also possible that the calibrated parameters are unable to capture behaviors that were not exhibited within the experimental datasets used in parameterization. While we tested ranges for each parameter and settled on a single parameter set that best fit the calibration data, there may be additional parameters sets that fit the calibration data but have varied levels of stochasticity and altered reproducibility of replicate simulations. Lastly, the current model was calibrated to male mice data despite known sex difference in skeletal muscle, regeneration mechanisms, and the timeline of recovery^128–130^. Experimental measurements of female muscle regeneration are fairly limited because most muscle injury studies only use male mice or do not distinguish between sexes, making it difficult to incorporate sex differences into the model^131^. Experiments that incorporate female mice and measure hormone levels are needed to accurately incorporate rules to distinguish between the sex dependent dynamics of muscle regeneration.

This paper describes a significant advancement in modeling the complex process of muscle regeneration. Future efforts will extend the use of parameter density estimation to optimize the selection, doses, and timing of injections of exogenously delivered cytokines. Further refinement of analysis methods could be pursued to disentangle specific underlying mechanisms of the dynamic feedbacks that drive the observed model outputs. Predictions from model simulations will also be used to inform future experiments by highlighting crucial time points to measure and predicted effect sizes for power analysis. Additionally, we aim to explore diverse muscle injury types and locations (i.e. injury relative to microvascular components) and their varying recovery responses, addressing challenges in comparing different acute injury techniques found in the literature. This study underscores the significance of cellular and cytokine spatial dynamics in muscle regeneration. Further inclusion of additional factors and hormones would provide a more holistic understanding of the system and how treatments may be altered based on microenvironmental conditions, providing a unique framework for the study of personalized muscle injury treatment.

## Acknowledgements

The authors acknowledge NIH Grant #R21AR080415, Wu-Tsai Foundation Agility Project Funding, and NSF GRFP GRANT #1842490 for financially supporting this research and to James Glazier and T.J. Sego for providing technical support with the CC3D ABM platform.

## Additional Resources

The ABM source code is publicly available at the following sites:

SimTK: https://simtk.org/docman/?group_id=2635

Zendo: https://zenodo.org/records/10403014

GitHub: https://github.com/mh2uk/ABM-of-Muscle-Regeneration-with-Microvascular-Remodeling

## Supplemental

**Supplemental Figure 1.**
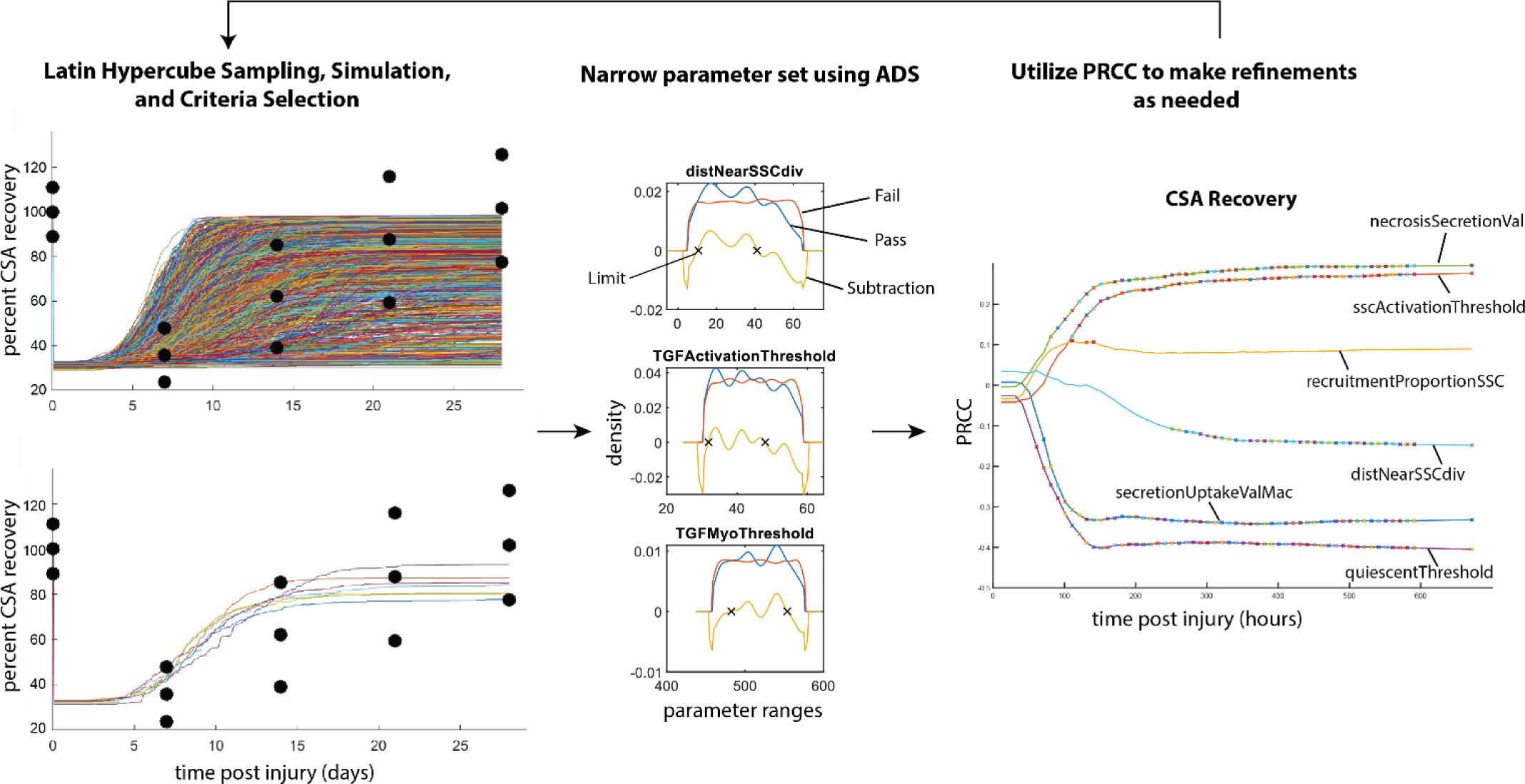
Overview of Calibration Methods. Latin hypercube sampling is used to generate 600 unique parameter sets given starting bounds, each of which was run in triplicate. The simulations were filtered given specified criteria (i.e. fitting within experimental bounds for CSA recovery) and then alternative density subtraction (ADS) was used to narrow in the parameter bounds. Partial rank squared correlation coefficient (PRCC) was used to gain insight into model sensitivity and adjust the bounds in case initial parameter bounds were too wide or too narrow. This method also allowed for model rule execution refinement to correct cases that interfere with the dynamics of other cell types.

**Supplemental Table 1.** Unknown model parameters calibrated using LHS to recapitulate published literature.

| Parameter | Description | Range Tested | Value |
| --- | --- | --- | --- |
| neutroRecruitmentProportion | Number of neutrophils generated at every recruitment as proportion of necrotic cells | (4, 12) | 7.8322 |
| secretionUptakeValNeu | Cytokine secreted by neutrophil per step during and after "eating necrosis" also max uptake value | (20, 80) | 37.3001 |
| necrosisThreshold | Neutrophils are recruited as long as the necrosis amount is greater than this parameter | (0.75, 0.98) | 0.9202 |
| macroRecruitmentProportion | Recruitment of macrophages is proportional to mean MCP-1 | (0.1, 0.7) | 0.4701 |
| secretionUptakeValMac | Cytokine secreted by macrophages per step, also max uptake value | (0.3, 95) | 48.2090 |
| TNFconverter | Value of TNF-α to convert monocyte to M1 | (15, 150) | 109.9025 |
| macRecruitThres | Required mean MCP-1 to signal recruitment of macrophage | (0.05, 4.5) | 1.4289 |
| macRecruitStop | Required mean TGF-β to signal stop of macrophage recruitment | (0.05, 0.4) | 0.1406 |
| IL10TransThreshold | Local IL-10 that can induce M1 to M2 transition | (15, 600) | 78.5442 |
| phagoThresholdforNeuM2 | Phagocytosis threshold for neutrophil apoptosis or M1 to M2 transition | (0, 8) | 2.0706 |
| sscActivationThreshold | Level of HGF to activate SSC | (10, 100) | 44.7091 |
| recruitmentProportionSSC | Number of SSC recruited every time step is proportional to sum of HGF + MMP-9 – TGF-β means | (0.08, 1) | 0.8213 |
| quiescentProb | Probability of SSC apoptosis vs remaining quiescent | (1, 4) | 3.3617 |
| secretionUptakeValSSC | Cytokine secreted by SSC per step when activated also max uptake value | (0.5, 200) | 44.4650 |
| sscTGFApoptosisThreshold | Amount of TGF-β to induce apoptosis | (50, 190) | 79.1563 |
| sscApoptosisProb | Probability of SSC apoptosis with apoptosis signal | (0.2, 0.99) | 0.7651 |
| VEGFblockApop | Amount of VEGF-A required to block SSC apoptosis | (60, 190) | 80.6055 |
| SSCdivisionThreshold | TNF-α + VEGF-A – TGF-β has to be greater than this for SSC to divide | (50, 400) | 126.6268 |
| SSCdiffThreshold | IL-10 – HGF – TNF-α – TGF-β has to be greater than this for SSC diff | (5, 145) | 57.9870 |
| sscDivideProb | Probability of SSC to divide if signal is not present | (0.01, 0.6) | 0.0972 |
| sscDiffProb | Probability for SSC to differentiate without signal | (0.06, 1) | 0.6136 |
| fuseProb | Probability of SSC fusing even if the collagen amount is still low | (0.06, 1) | 0.3 |
| quiescentThreshold | SSC will return to quiescence if HGF is lower than this at their location | (0.2, 100) | 36.3109 |
| necrosisSecretionVal | Amount of HGF and TGF-β secreted by necrotic cells per step | (0.0005, 35) | 21.9514 |
| VEGFsecretionValFib | VEGF-A secreted by fibers per step | (0.02, 0.1) | 0.0498 |
| VEGF_MMPthreshold | Level of VEGF-A to trigger sprouting/elongation from non-perfused capillaries | (98, 800) | 469.0449 |
| capVEGFUptakeMax | Max uptake of VEGF-A by capillaries | (8, 32) | 23.8174 |
| MMPsecretionValCap | MMP-9 secreted as capillary is regenerating/sprouting | (4, 16) | 8.2444 |
| NumFibers | Number of fibers nearby must be higher than this to check for capillary distances | (1, 2) | 1.3899 |
| sproutProb | Probability of capillary sprouting behavior | (0.08, 0.32) | 0.1566 |
| secretionUptakeValFibro | Cytokine secretion from fibroblasts per mcs also max uptake value | (0.3, 77) | 9.0436 |
| collagenSecretionValFibro | Collagen secretion from fibroblasts per timestep when it is above ECM | (0.09, 0.24) | 0.1388 |
| distNearSSCdiv | If fibroblast is within this distance of a dividing SSC, it will divide | (5, 65) | 35 |
| TGFActivationThreshold | Level of TGF-β required to activate fibroblast | (20, 80) | 40.3643 |
| TGFMyoThreshold | Level of sustained TGF-β required to turn fibroblast into myofibroblast | (280, 650) | 536.7525 |
| fibroblastTNFApoptosisThreshold | Level of TNF-α to initiate fibroblast apoptosis | (100, 550) | 310.5323 |
| TGFBlockApoptosisThreshold | Level of TGF-β required to block TNF-initiated apoptosis | (160, 1000) | 848.3706 |
| fibroblastApoptosisProb | probability of fibroblast apoptosis if TNF-α and TGF-β conditions are met | (0.65, .99) | 0.8423 |
| IL10decay | Decay rate of IL-10 | (0.00046, 0.00184) | 0.0012 |
| HGFdecay | Decay rate of HGF | (0.000135, 0.0003) | 0.000228 |
| TGFdecay | Decay rate of TGF | (0.0005, 0.05) | 0.0242 |
| TNFdecay | Decay rate of TNF | (0.0000682, 0.002260) | 0.00076272 |
| MCPdecay | Decay rate of MCP-1 | (0.00596, 0.02384) | 0.0124 |
| MMPdecay | Decay rate of MMP-9 | (0.0012, 0.1) | 0.0112 |
| VEGFdecay | Decay rate of VEGF-A | (0.000464, 0.005569) | 0.0015 |
| MinCapDist | Minimum distance between capillary and another capillary | (2.5, 40) | 9.6060 |
| capPlacementThreshold | Threshold for if a capillary will be placed/distance required between fiber and cap | (8, 45) | 34.6346 |
| M2prolifProb | Probability of M2 proliferation behavior occurring | (0.145305, 0.60) | 0.2 |
| M1HalfLife | Corresponds with rate of M1 apoptosis | (15, 55) | 24.6360 |
| M2HalfLife | Corresponds with rate of M2 apoptosis | (30, 70) | 58.1243 |
| reperfusionLag | Probability of capillary regaining perfusion | (0.001, 0.98) | 0.1 |

**Supplemental Table 2.** Criteria utilized for CaliPro model calibration.

| Iteration | Criteria |
| --- | --- |
| 1 | End criteria: SSC, M1 macrophages, M2 macrophage, fibroblasts, and neutrophils must be below max threshold at the final timepoint (threshold held constant for all iterations). |
| 2 | SSC, CSA, and end criteria: within 4 times the experimental std for SSC and 1 times CSA std. |
| 3 | Fibroblast, SSC, CSA, replicate check, and end criteria: within 2.1 times the experimental std for fibroblasts, 2.5 for SSC, and 1 times CSA std. More than 1 replicate of the parameter set must meet the criteria. |
| 4 | Fibroblast, SSC, CSA, and end criteria: within 2.2 times the experimental std for fibroblasts and SSC and 1 times CSA std. |
| 5 | Fibroblast, SSC, CSA, and end criteria: within 1.5 times the experimental std for fibroblasts, 1.75 for SSC, and 1 times CSA std. |
| 6 | Fibroblast, SSC, CSA, and end criteria: within 1.75 times the experimental std for fibroblasts, 1.5 for SSC, and 1 times CSA std. |
| 7 | Fibroblast, SSC, CSA, replication, and end criteria: within 2.5 times the experimental std for fibroblasts and SSC, and 1 times CSA std. All replicates are required to pass for the parameter set to pass. |

**Supplemental Figure 2.**
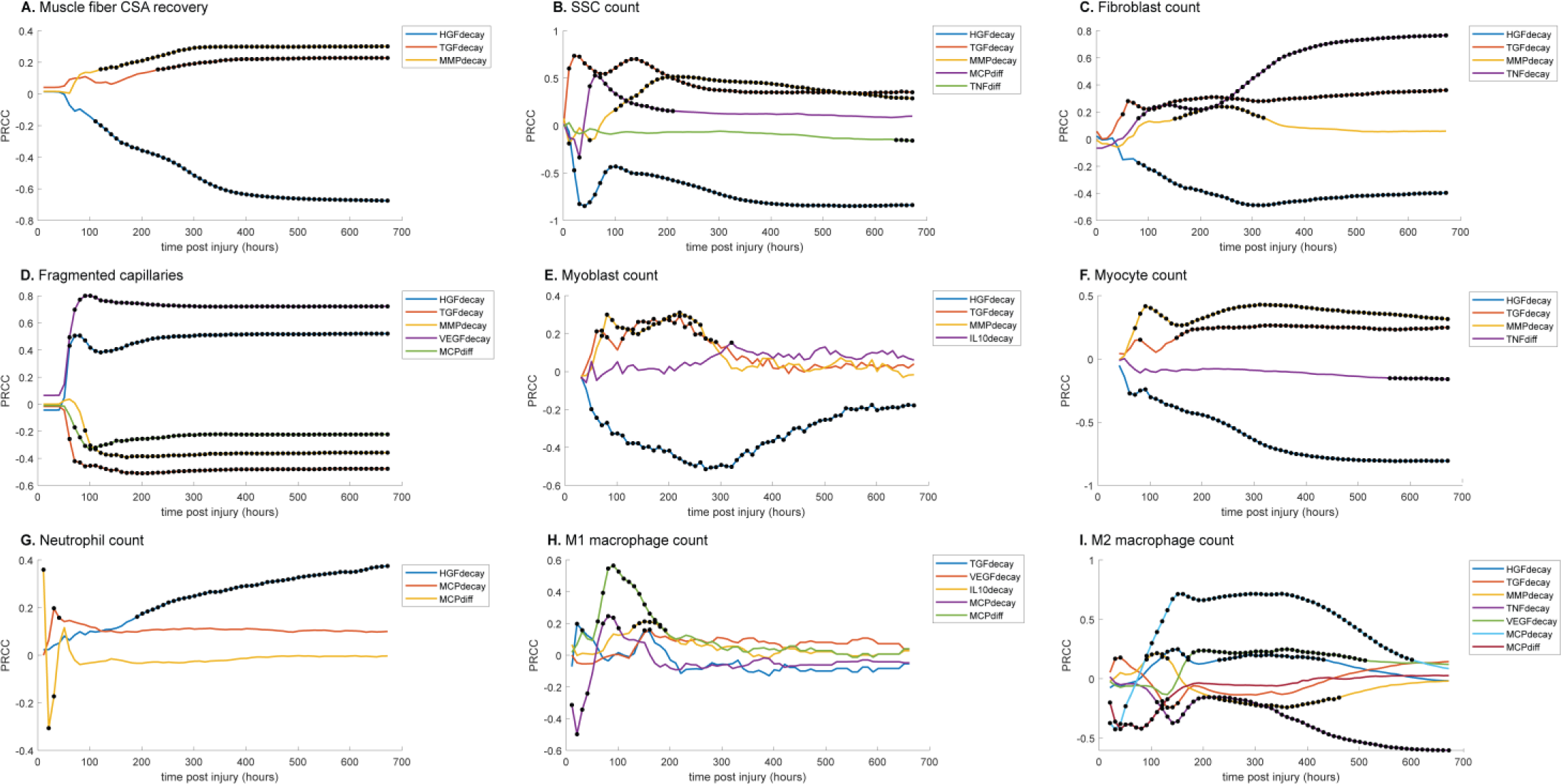
PRCC plots for various model outputs over time to illustrate how the significance of cytokine decay and diffusion parameters varies at different points throughout regeneration. Black dots indicate statistically significant (P < 0.05) correlation for that timepoint. (A) CSA recovery had correlations with HGF, TGF-β, and MMP-9 decay. (B) SSC count was correlated with HGF, TGF-β, MMP-9 decay and MCP-1 and TND diffusion. (C) Fibroblast count was correlated with HGF, TGF-β, MMP-9, and TNF-α decay. (D) HGF, TGF-β, MMP-9, VGEF decay and MCP-1 diffusion were correlated with the number of non-perfused capillaries. (E) Myoblast cell count was correlated with HGF, TGF-β, MMP-9, and IL-10 decay. (F) Myocyte cell count was correlated with HGF, TGF-β, and MMP-9 decay and TNF-α diffusion. (G) HGF and MCP-1 decay as well as MCP-1 diffusion were correlated with neutrophil count. (H) M1 macrophage cell count was correlated with TGF-β, VEGF-A, IL-10, and MCP-1 decay and MCP-1 diffusion. (I) M2 macrophage count was correlated with HGF, TGF-β, MMP-9, TNF-α, VEGF-A, MCP-1 decay and MCP-1 diffusion.

**Supplemental Figure 3.**
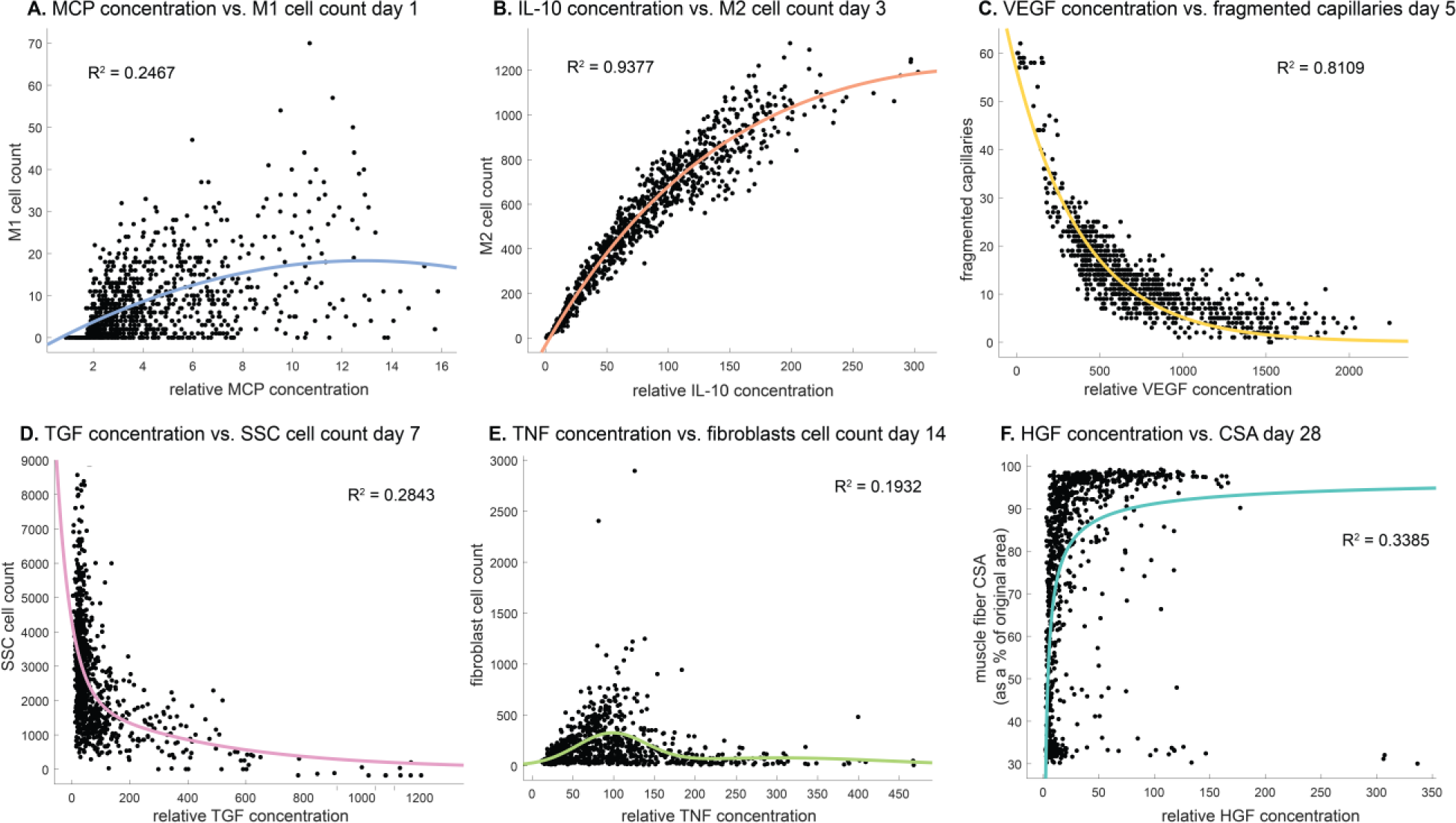
Cytokine concentrations are correlated with cell counts and recovery metrics at various stages of regeneration. There is an optimal MCP-1 concentration that tends to result in higher M1 counts 1 day post injury (A). IL-10 concentration is positively correlated with M2 count 3 days post injury (B). VEGF-A concentration is negatively correlated with the number of fragmented (non-perfused) capillaries 5 days post injury (C). Higher TGF-β concentrations tends to result in lower SSC cell count 7 days post injury (D). Fibroblasts cell count is highest at an optimal TNF-α concentration with higher or lower levels hindering cell count 14 days post injury (E). HGF concentration is positively correlated with CSA recovery at day 28 post injury but there appears to be a threshold where high HGF is no longer correlated with increased recovery (F).

**Supplemental Table 3.**
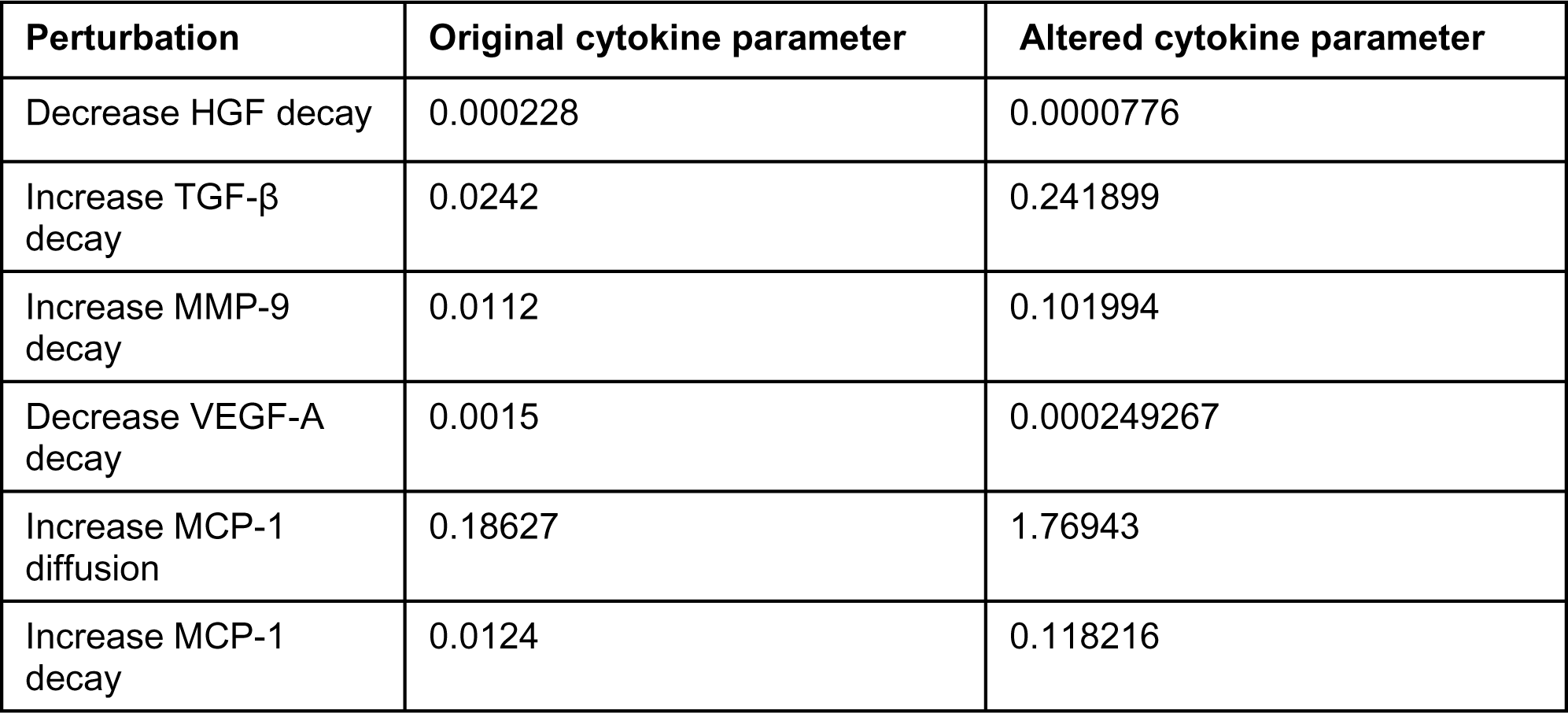
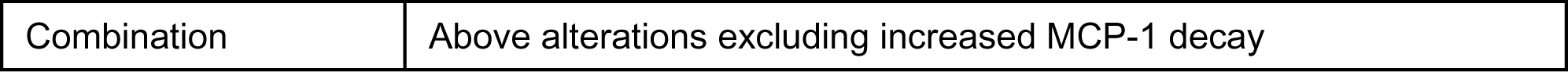
Cytokine perturbations based on PRCC.

| Perturbation | Original cytokine parameter | Altered cytokine parameter |
| --- | --- | --- |
| Decrease HGF decay | 0.000228 | 0.0000776 |
| Increase TGF-β decay | 0.0242 | 0.241899 |
| Increase MMP-9 decay | 0.0112 | 0.101994 |
| Decrease VEGF-A decay | 0.0015 | 0.000249267 |
| Increase MCP-1 diffusion | 0.18627 | 1.76943 |
| Increase MCP-1 decay | 0.0124 | 0.118216 |

|  |  |
| --- | --- |
| Combination | Above alterations excluding increased MCP-1 decay |

**Supplemental Figure 4.**
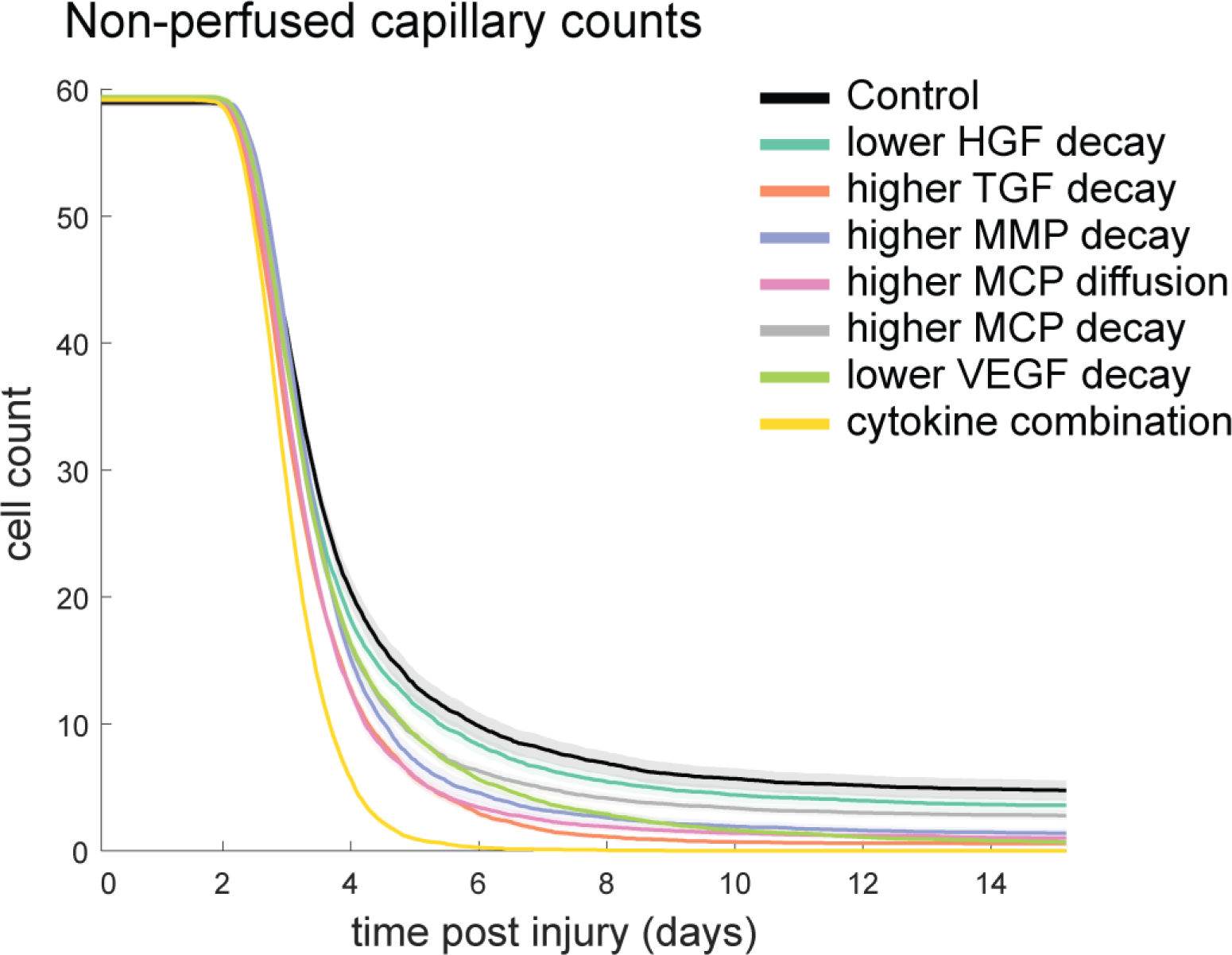
Non-perfused capillaries for each cytokine perturbation. The combined cytokine perturbation had the lowest number of non-perfused capillaries and all other perturbations resulted in less non-perfused capillaries compared to the control.

**Supplemental Figure 5.**
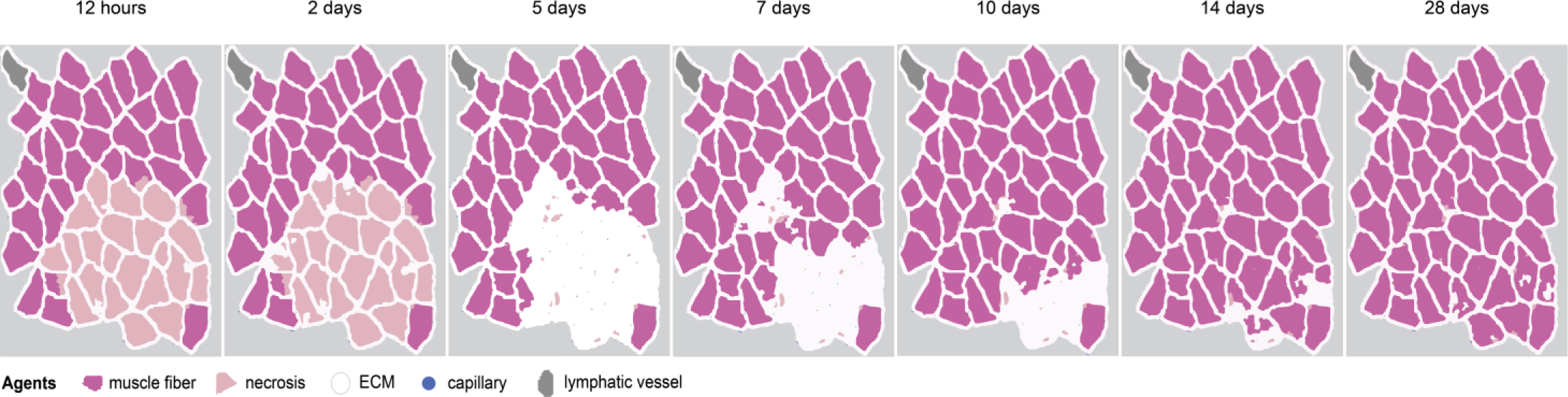
Overview of ABM simulation with different initial histology configuration.

## Supplemental Text 1. Cellular-Potts

### Cellular-Potts model implementation

In the CPM, individual cells are represented as a collection of pixels on a square lattice. These computational representations of cells have properties of predefined volume, contact energy with surrounding cells in their environment, and affinity or aversion to diffusing species that drive chemotactic behaviors^23^. These properties are defined mathematically in the Effective Energy function *H* which is evaluated in every computational timestep. Equation (1) represents the effective energy function that imposes the physical properties of all cells in the agent-based CPM.

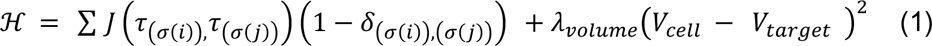

The first term describes contact energy of each cell for neighboring cells in its environment governed by *J* the contact coefficient, where *i,j* denote neighboring lattice sites, τ denotes cell types, and σ denotes individual cells in the simulation. The delta function localizes contact energy contributions to cell-cell interfaces. The second term represents a volume constraint scaled by *λ_volume_* on each cell. Motile cells in the simulation that exhibit chemotactic behavior towards a target have additional terms to drive their movement described in (2).

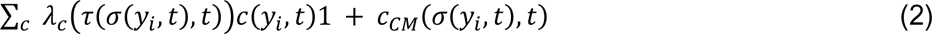

Which models logarithmic chemotaxis by cell type and chemokine field concentration influenced by chemotaxis parameter *λ*_*c*_, chemical field concentration *c*, and cell body center-of-mass measurement *c_CM_* from which chemotaxis behaviors are calculated. Diffusion and secretion behavior of chemical species is governed by the general diffusion equation described in (3).

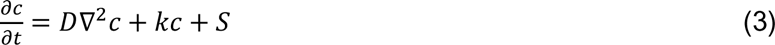

Where *k* defines the decay constant of diffusive species concentration *c*, for which *D* is the diffusion constant. The *S* term is an additional term to represent secretion of diffusive species.

To apply these relations to recreate cell movement, the CPM randomly selects a pair of neighboring pixels and evaluates whether one pixel may copy itself to the location of the other, called a “copy attempt.” The probability of acceptance or rejection of that pixel copy attempt is governed described by the Boltzmann acceptance function described in (4).

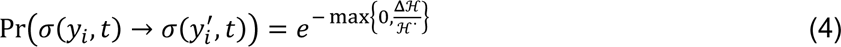

This function describes the probability of the copy attempt 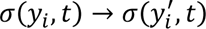 where pixel 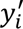 copies to pixel *y_i_* in the CPM algorithm^23^. The value ℋ* represents our *temperature* parameter in the CPM, or the parameter that affects stochasticity of lattice site copy attempts. Δℋ is the change in effective energy due to a lattice site copy attempt. The Cellular-Potts simulation discussed here was implemented using CompuCell3D (CC3D), an open-source Python-based CPM platform^23^.

### Cellular-Potts model initialization

The muscle cross-section geometry was initialized by importing a histology image of (murine gastrocnemius stained with laminin α2) masked to distinguish between muscle fiber and ECM. The mask was imported into an initialization CC3D script that defined the muscle fibers, ECM, and microvasculature to specific Cellular-Potts lattice types and generated a Potts Initialization File (PIF) that was imported into the ABM as the starting cross-section. This initial configuration resulted in a square lattice of dimensions 321×417 pixels. The model consisted of 2 layers in the z axis: a tissue environment layer consisting of muscle, ECM, capillaries, and lymphatic channels, and an immune cell layer where immune cell agents migrated and interacted in the “2.5D” model. CPM interaction neighbor order was 4 for typical adhesion energy terms, and 1 for all other CPM Hamiltonian terms. Cellular-Potts model parameters are documented in Supplemental Table 4, adhesion terms are described in Supplemental Table 5. Non-zero parameter values for adhesion energies between lattice types were calibrated visually to prevent aggregation of migratory cell types.

**Supplemental Table 4.** CPM model parameters.

| Parameter | Value | Justification |
| --- | --- | --- |
| Simulation lattice size | 321x417x2 | 2 z-layers: Tissue microenvironment layer and migratory cell layer |
| Metropolis Algorithm Temperature | 10 | This is a default, numerically stable temperature for CPM models described in CompuCell3D |
| Neighbor Order | 1 | Interactions spanning 1 pixel distance for Cellular-Potts cell agents was sufficient for agent-based behaviors |

**Supplemental Table 5.** CPM agent Adhesion parameters.

| CPM Lattice Interaction | Parameter Value |
| --- | --- |
| Medium - Medium | 0 |
| Medium - Fiber | 0 |
| Medium - ECM | 0 |
| Medium - SSC | 0 |
| Medium - Capillary | 0 |
| Medium - Neutrophil | 0 |
| Medium - Macrophage | 0 |
| Medium - Necrotic | 0 |
| Medium - Wall | 0 |
| Medium - Lymphatic | 0 |
| Fiber - Fiber | 0 |
| Fiber - ECM | 0 |
| Fiber - SSC | 35 |
| Fiber - Capillary | 0 |
| Fiber - Neutrophil | 35 |
| Fiber - Macrophage | 20 |
| Fiber - Necrotic | 0 |
| Fiber - Wall | 0 |
| Fiber - Lymphatic | 0 |
| ECM - ECM | 0 |
| ECM - SSC | 30 |
| ECM - Capillary | 0 |
| ECM - Neutrophil | 30 |
| ECM - Macrophage | 20 |
| ECM - Necrotic | 0 |
| ECM - Wall | 0 |
| ECM - Lymphatic | 0 |
| SSC - SSC | 35 |
| SSC - Capillary | 30 |
| SSC - Neutrophil | 15 |
| SSC - Macrophage | 15 |
| SSC - Necrotic | 30 |
| SSC - Wall | 20 |
| SSC - Lymphatic | 10 |
| Capillary - Capillary | 0 |
| Capillary - Neutrophil | 35 |
| Capillary - Macrophage | 20 |
| Capillary - Necrotic | 0 |
| Capillary - Wall | 0 |
| Capillary - Lymphatic | 0 |
| Neutrophil - Neutrophil | 35 |
| Neutrophil - Macrophage | 15 |
| Neutrophil - Necrotic | 35 |
| Neutrophil - Wall | 20 |
| Neutrophil - Lymphatic | 10 |
| Macrophage - Macrophage | 15 |
| Macrophage - Necrotic | 20 |
| Macrophage - Wall | 20 |
| Macrophage - Lymphatic | 10 |
| Necrotic - Necrotic | 0 |
| Necrotic - Wall | 0 |
| Necrotic - Lymphatic | 0 |
| Wall - Wall | 0 |
| Wall - Lymphatic | 0 |
| Lymphatic - Lymphatic | 0 |
| Fibroblast - Medium | 0 |
| Fibroblast - Fiber | 20 |
| Fibroblast - ECM | 20 |
| Fibroblast - SSC | 15 |
| Fibroblast - Capillary | 20 |
| Fibroblast - Neutrophil | 15 |
| Fibroblast - Macrophage | 15 |
| Fibroblast - Necrotic | 20 |
| Fibroblast - Wall | 20 |
| Fibroblast - Lymphatic | 10 |
| Fibroblast - Fibroblast | 15 |
| Neighbor Order | 4 |

**Supplemental Table 6.** Experimental data description for model comparison.

| Experimental data description and relevant figure from literature | Model comparison |
| --- | --- |
| CTX injury to mouse hindlimb muscle and quantification of average cross-sectional area ($\mu\text{m}^2$) of individual muscle fibers for a given animal was determined after manually outlining individual muscle fibers in each of the digitally captured images (Fig. 3B)<sup>102</sup> | Muscle CSA recovery calibration (Fig. 3A) |
| BaCl<sub>2</sub> injury to mouse hindlimb and IF quantification of SSC (Pax7+) and fibroblasts (Tcf4+) cells (Fig 2DD & EE)<sup>76</sup> | SSC and fibroblast count calibration (Fig. 3B & C) |
| NTX injury to mouse hindlimb and macrophage quantification based on number of F4/80+ cells (Fig. 4E)<sup>97</sup> | Total macrophage count validation (Fig. 3D) |
| CTX injury to mouse hindlimb muscle and capillary density was calculated using biotinylated lectin and H&E (Fig. 6B)<sup>102</sup> | Capillaries/$\text{mm}^2$ validation (Fig. 3H) |
| BaCl<sub>2</sub> injury to mouse hindlimb and flow cytometry to assess CD45<sup>+</sup>/F4/80<sup>+</sup>/Ly6C<sup>hi</sup> (M1) and CD45<sup>+</sup>/F4/80<sup>+</sup>/Ly6C<sup>lo</sup> (M2) macrophages (Fig. 2A)<sup>107</sup> | M1 and M2 cell count validation (Fig. 3E & H) |
| CTX injury to mouse hindlimb muscle and IHC analysis was used to label neutrophils using Ly6G antibody (Fig. 3A)<sup>108</sup> | Neutrophil cell count validation (Fig. 3G) |
| Bothrops asper snake venom induced hindlimb injury to mice pretreated with either an antimouse granulocyte rat monoclonal immunoglobulin G (IgG) antibody or with isotype-matched control antibody. Inflammatory infiltrate was quantified from muscle tissue (Fig. 7A) and the extent of regeneration was quantitatively assessed by determining the percentage of the area comprised of regenerating muscle fibers in relation to the total area of muscle damage<sup>109</sup> | Neutrophil depletion perturbation (Fig. 4) |
| IL-4-conjugated gold nanoparticles were injected into murine skeletal muscle following hindlimb ischemia. M1 and M2 cells were quantified with flow cytometry (Fig. 6 A, C, and E) and muscle force was measured for quantification of regeneration (Fig. 4G)<sup>164</sup> | Anti-inflammatory nanoparticles perturbation (Fig. 4) |
| IL-10 null mice underwent hindlimb unloading and reloading to induce muscle damage. Muscle CSA (Fig. 6C), M1 and M2 cells (Fig. 3A-B), and SSC differentiation markers were quantified (Fig. 3C-F)<sup>163</sup> | IL-10 KO perturbation (Fig. 4) |
| Noetoxin or freeze injury to CXCL12Gagtm/Gagtm and control mouse hindlimb muscle with H&E for assessment of muscle regeneration (Fig. 3), transcriptome analysis to assess collagen organization and synthesis (Fig. 2C), and immune cell infiltration (Fig. 4S)<sup>111</sup> | Hindered angiogenesis perturbation (Fig. 4) |
| Following CTX injury to mouse hindlimb muscle and injection of intramuscular administration of rAAV vectors injury area was quantified for various doses of AAV-VEGF (Fig. 3D) and H&E was used to assess muscle recovery (Fig. 5)<sup>73</sup> | VEGF-A injection perturbation (Fig. 4) |
| CCL2<sup>-/-</sup> and control mice received BaCl<sub>2</sub> injection to induce acute injury. CSA quantification was used to assess muscle regeneration (Fig. 1I). Flow cytometry was used to quantify macrophage (Fig. 2A), and neutrophil (Fig. 2B) cells<sup>115</sup> | MCP-1 KO perturbation (Fig. 4) |
| CTX induced injury to control and TNF-α receptor double-knockout mice. Force was measured to quantify muscle recovery (Fig. 7), immunohistochemical staining of Mac-1 for quantifying immune cells (Fig. 6B), and western blot analysis was used to assess SSC differentiation (Fig. 2)<sup>165</sup> | TNF-α KO perturbation (Fig. 4) |
| Mice were treated with clodronate-containing or control liposomes and underwent a contusion injury. H&E was used for muscle regeneration assessment (Fig. 2), HGF (Fig. 4A) and TGF (Fig. 5A) mRNA were quantified<sup>110</sup> | Macrophage depletion perturbation (Fig. 4) |

## References

1. Quintero AJ, Wright VJ, Fu FH, Huard J. Stem Cells for the Treatment of Skeletal Muscle Injury Sports injury Stem cells Tissue engineering Fibrosis Regeneration Skeletal muscle. Clin Sports Med. 2009;28:1–11. doi:10.1016/j.csm.2008.08.009

2. Barroso GC, Thiele ES. MUSCLE INJURIES IN ATHLETES. Rev Bras Ortop (English Ed. 2011;46(4):354–358. doi:10.1016/s2255-4971(15)30245-7

3. Valle X. Clinical practice guide for muscular injuries: epidemiology, diagnosis, treatment and prevention. Br J Sports Med. 2011;45(2):e2–e2. doi:10.1136/bjsm.2010.081570.20

4. Järvinen TAH, Järvinen TLN, Kääriäinen M, et al. Muscle injuries: optimising recovery. Best Pract Res Clin Rheumatol. 2007;21(2):317–331. doi:10.1016/j.berh.2006.12.004

5. Huard J, Li Y, Fu FH. Muscle injuries and repair: Current trends in research. J Bone Jt Surg - Ser A. Published online 2002. doi:10.2106/00004623-200205000-00022

6. Forcina L, Cosentino M, Musarò A. Mechanisms Regulating Muscle Regeneration: Insights into the Interrelated and Time-Dependent Phases of Tissue Healing. Cells. 2020;9(5). doi:10.3390/cells9051297

7. Howard EE, Pasiakos SM, Blesso CN, Fussell MA, Rodriguez NR. Divergent Roles of Inflammation in Skeletal Muscle Recovery From Injury. Front Physiol. 2020;11:87. doi:10.3389/fphys.2020.00087

8. Westman AM, Peirce SM, Christ GJ, Blemker SS. Agent-based model provides insight into the mechanisms behind failed regeneration following volumetric muscle loss injury. Merks RMH, ed. PLOS Comput Biol. 2021;17(5):e1008937. doi:10.1371/journal.pcbi.1008937

9. Wagatsuma A. Endogenous expression of angiogenesis-related factors in response to muscle injury. Mol Cell Biochem 2006 2981. 2006;298(1):151-159. doi:10.1007/S11010-006-9361-X

10. Chen CH, Chang CH, Lee CH. Usage of Growth Factors in Acute Muscle Injuries. In: Sports Injuries. ; 2015:2343–2351. doi:10.1007/978-3-642-36569-0_174

11. Husmann I, Soulet L, Gautron J, Martelly I, Barritault D. Growth factors in skeletal muscle regeneration. Cytokine Growth Factor Rev. 1996;7(3):249–258. doi:10.1016/S1359-6101(96)00029-9

12. Itoh Y. Proteolytic modulation of tumor microenvironment signals during cancer progression. Front Oncol. 2022;12. doi:10.3389/FONC.2022.935231

13. Ciano-Petersen NL, Muñiz-Castrillo S, Birzu C, et al. Cytokine dynamics and targeted immunotherapies in autoimmune encephalitis. Brain Commun. 2022;4(4):1–16. doi:10.1093/braincomms/fcac196

14. Ferrara N. Binding to the Extracellular Matrix and Proteolytic Processing: Two Key Mechanisms Regulating Vascular Endothelial Growth Factor Action. Mol Biol Cell. 2010;21(5):687. doi:10.1091/MBC.E09-07-0590

15. Virgilio KM, Martin KS, Peirce SM, Blemker SS. Agent-based model illustrates the role of the microenvironment in regeneration in healthy and mdx skeletal muscle. J Appl Physiol. 2018;125(5):1424–1439. doi:10.1152/japplphysiol.00379.2018

16. Martin KS, Virgilio KM, Peirce SM, Blemker SS. Computational modeling of muscle regeneration and adaptation to advance muscle tissue regeneration strategies. Cells Tissues Organs. 2016;202(3-4):250–266. doi:10.1159/000443635

17. Khuu S, Fernandez JW, Handsfield GG. A Coupled Mechanobiological Model of Muscle Regeneration In Cerebral Palsy. Front Bioeng Biotechnol. 2021;9. doi:10.3389/fbioe.2021.689714

18. Khuu S, Fernandez JW, Handsfield GG. Delayed skeletal muscle repair following inflammatory damage in simulated agent-based models of muscle regeneration. Peirce SM, ed. PLOS Comput Biol. 2023;19(4):e1011042. doi:10.1371/JOURNAL.PCBI.1011042

19. Virgilio KM, Jones BK, Miller EY, et al. Computational Models Provide Insight into In Vivo Studies and Reveal the Complex Role of Fibrosis in mdx Muscle Regeneration. Ann Biomed Eng. Published online 2020. doi:10.1007/s10439-020-02566-1

20. Joslyn LR, Kirschner DE, Linderman JJ. CaliPro: A Calibration Protocol That Utilizes Parameter Density Estimation to Explore Parameter Space and Calibrate Complex Biological Models. Cell Mol Bioeng. 2021;14(1):31–47. doi:10.1007/s12195-020-00650-z

21. Martin KS, Blemker SS, Peirce SM. Agent-based computational model investigates muscle-specific responses to disuse-induced atrophy. J Appl Physiol. 2015;118(10):1299–1309. doi:10.1152/japplphysiol.01150.2014

22. Gianlupi JF, Mapder T, Sego TJ, et al. Multiscale Model of Antiviral Timing, Potency, and Heterogeneity Effects on an Epithelial Tissue Patch Infected by SARS-CoV-2. Viruses 2022, Vol 14, Page 605. 2022;14(3):605. doi:10.3390/V14030605

23. Swat MH, Thomas GL, Belmonte JM, Shirinifard A, Hmeljak D, Glazier JA. Multi-Scale Modeling of Tissues Using CompuCell3D. Methods Cell Biol. 2012;110:325. doi:10.1016/B978-0-12-388403-9.00013-8

24. Stephenson ER, Kojouharov H V. A mathematical model of skeletal muscle regeneration. Math Methods Appl Sci. 2018;41(18):8589–8602. doi:10.1002/mma.4908

25. Waldemer-Streyer RJ, Kim D, Chen J. Muscle cell-derived cytokines in skeletal muscle regeneration. FEBS J. 2022;289(21):6463–6483. doi:10.1111/febs.16372

26. Rucavado A, Escalante T, Teixeira CFP, Fernándes CM, Díaz C, Gutiérrez JM. Increments in cytokines and matrix metalloproteinases in skeletal muscle after injection of tissue-damaging toxins from the venom of the snake Bothrops asper. Mediators Inflamm. 2002;11(2):121–128. doi:10.1080/09629350220131980

27. Yin H, Price F, Rudnicki MA. Satellite cells and the muscle stem cell niche. Physiol Rev. 2013;93(1):23–67. doi:10.1152/PHYSREV.00043.2011

28. Madaro L, Bouché M. From innate to adaptive immune response in muscular dystrophies and skeletal muscle regeneration: The role of lymphocytes. Biomed Res Int. 2014;2014. doi:10.1155/2014/438675

29. Wang X, Hossain M, Bogoslowski A, Kubes P, Irimia D. Chemotaxing neutrophils enter alternate branches at capillary bifurcations. Nat Commun 2020 111. 2020;11(1):1-12. doi:10.1038/s41467-020-15476-6

30. Butterfield TA, Best TM, Merrick MA. The dual roles of neutrophils and macrophages in inflammation: A critical balance between tissue damage and repair. J Athl Train. 2006;41(4):457–465. doi:10.1016/s0162-0908(08)79217-1

31. Wang J. Neutrophils in tissue injury and repair. Cell Tissue Res. 2018;371(3):531–539. doi:10.1007/s00441-017-2785-7

32. Soehnlein O, Zernecke A, Eriksson EE, et al. Neutrophil secretion products pave the way for inflammatory monocytes. Blood. 2008;112(4):1461. doi:10.1182/BLOOD-2008-02-139634

33. Fox S, Leitch AE, Duffin R, Haslett C, Rossi AG. Neutrophil Apoptosis: Relevance to the Innate Immune Response and Inflammatory Disease. J Innate Immun. 2010;2(3):216. doi:10.1159/000284367

34. Oishi Y, Manabe I. Macrophages in inflammation, repair and regeneration. Int Immunol. 2018;30(11):511–528. doi:10.1093/INTIMM/DXY054

35. Elkington PT, Green JA, Friedland JS. Analysis of matrix metalloproteinase secretion by macrophages. Methods Mol Biol. 2009;531:253–265. doi:10.1007/978-1-59745-396-7_16

36. Chen GY, Nuñez G. Sterile inflammation: sensing and reacting to damage. Nat Rev Immunol 2010 1012. 2010;10(12):826-837. doi:10.1038/nri2873

37. Lacy P, Stow JL. Cytokine release from innate immune cells: association with diverse membrane trafficking pathways. Blood. 2011;118(1):9–18. doi:10.1182/BLOOD-2010-08-265892

38. Vogel DYS, Heijnen PDAM, Breur M, et al. Macrophages migrate in an activation-dependent manner to chemokines involved in neuroinflammation. J Neuroinflammation. 2014;11:23. doi:10.1186/1742-2094-11-23

39. Molnarfi N, Benkhoucha M, Funakoshi H, Nakamura T, Lalive PH. Hepatocyte growth factor: A regulator of inflammation and autoimmunity. Autoimmun Rev. 2015;14(4):293–303. doi:10.1016/J.AUTREV.2014.11.013

40. Furrer R, Handschin C. Optimized engagement of macrophages and satellite cells in the repair and regeneration of exercised muscle. In: Research and Perspectives in Endocrine Interactions. Springer Verlag; 2017:57–66. doi:10.1007/978-3-319-72790-5_5

41. Kratofil RM, Kubes P, Deniset JF. Monocyte conversion during inflammation and injury. Arterioscler Thromb Vasc Biol. 2017;37(1):35–42. doi:10.1161/ATVBAHA.116.308198

42. Chazaud B, Sonnet C, Lafuste P, et al. Satellite cells attract monocytes and use macrophages as a support to escape apoptosis and enhance muscle growth. J Cell Biol. 2003;163(5):1133. doi:10.1083/JCB.200212046

43. Owen JL, Mohamadzadeh M. Macrophages and chemokines as mediators of angiogenesis. Front Physiol. 2013;4. doi:10.3389/FPHYS.2013.00159

44. Reibman J, Meixler S, Lee TC, et al. Transforming growth factor beta 1, a potent chemoattractant for human neutrophils, bypasses classic signal-transduction pathways. Proc Natl Acad Sci U S A. 1991;88(15):6805. doi:10.1073/PNAS.88.15.6805

45. Greenlee-Wacker MC. Clearance of apoptotic neutrophils and resolution of inflammation. Immunol Rev. 2016;273(1):357. doi:10.1111/IMR.12453

46. Watanabe S, Alexander M, Misharin A V., Budinger GRS. The role of macrophages in the resolution of inflammation. J Clin Invest. 2019;129(7):2619. doi:10.1172/JCI124615

47. Uribe-Querol E, Rosales C. Phagocytosis: Our Current Understanding of a Universal Biological Process. Front Immunol. 2020;11:1066. doi:10.3389/fimmu.2020.01066

48. Yoon YS, Lee YJ, Choi YH, Park YM, Kang JL. Macrophages programmed by apoptotic cells inhibit epithelial-mesenchymal transition in lung alveolar epithelial cells via PGE2, PGD2, and HGF. Sci Reports 2016 61. 2016;6(1):1–18. doi:10.1038/srep20992

49. D’Angelo F, Bernasconi E, Schäfer M, et al. Macrophages promote epithelial repair through hepatocyte growth factor secretion. Clin Exp Immunol. 2013;174(1):60. doi:10.1111/CEI.12157

50. Popov Y, Sverdlov DY, Bhaskar KR, et al. Macrophage-mediated phagocytosis of apoptotic cholangiocytes contributes to reversal of experimental biliary fibrosis. Am J Physiol - Gastrointest Liver Physiol. 2010;298(3):G323. doi:10.1152/AJPGI.00394.2009

51. Arnold L, Henry A, Poron F, et al. Inflammatory monocytes recruited after skeletal muscle injury switch into antiinflammatory macrophages to support myogenesis. J Exp Med. 2007;204(5):1057–1069. doi:10.1084/jem.20070075

52. Chung EY, Liu J, Homma Y, et al. Interleukin-10 Expression in Macrophages during Phagocytosis of Apoptotic Cells Is Mediated by the TALE homeoproteins Pbx-1 and Prep-1. Immunity. 2007;27(6):952. doi:10.1016/J.IMMUNI.2007.11.014

53. Mosser DM, Edwards JP. Exploring the full spectrum of macrophage activation. Nat Rev Immunol 2008 812. 2008;8(12):958-969. doi:10.1038/nri2448

54. Saini J, McPhee JS, Al-Dabbagh S, Stewart CE, Al-Shanti N. Regenerative function of immune system: Modulation of muscle stem cells. Ageing Res Rev. 2016;27:67–76. doi:10.1016/J.ARR.2016.03.006

55. Das A, Sinha M, Datta S, et al. Monocyte and Macrophage Plasticity in Tissue Repair and Regeneration. Am J Pathol. 2015;185(10):2596. doi:10.1016/J.AJPATH.2015.06.001

56. Arabpour M, Saghazadeh A, Rezaei N. Anti-inflammatory and M2 macrophage polarization-promoting effect of mesenchymal stem cell-derived exosomes. Int Immunopharmacol. 2021;97:107823. doi:10.1016/J.INTIMP.2021.107823

57. Reimann J, Irintchev A, Wernig A. Regenerative capacity and the number of satellite cells in soleus muscles of normal and mdx mice. Neuromuscul Disord. 2000;10(4-5):276–282. doi:10.1016/S0960-8966(99)00118-2

58. Kawamura K, Takano K, Suetsugu S, et al. N-WASP and WAVE2 acting downstream of phosphatidylinositol 3-kinase are required for myogenic cell migration induced by hepatocyte growth factor. J Biol Chem. 2004;279(52):54862–54871. doi:10.1074/JBC.M408057200

59. Wang W, Pan H, Murray K, Jefferson BS, Li Y. Matrix metalloproteinase-1 promotes muscle cell migration and differentiation. Am J Pathol. 2009;174(2):541–549. doi:10.2353/AJPATH.2009.080509

60. Allen RE, Boxhorn LK. Regulation of skeletal muscle satellite cell proliferation and differentiation by transforming growth factor-beta, insulin-like growth factor I, and fibroblast growth factor. J Cell Physiol. 1989;138(2):311–315. doi:10.1002/JCP.1041380213

61. González MN, de Mello W, Butler-Browne GS, et al. HGF potentiates extracellular matrix-driven migration of human myoblasts: Involvement of matrix metalloproteinases and MAPK/ERK pathway. Skelet Muscle. 2017;7(1):1–13. doi:10.1186/s13395-017-0138-6

62. Allen RE, Sheehan SM, Taylor RG, Kendall TL, Rice GM. Hepatocyte growth factor activates quiescent skeletal muscle satellite cells in vitro. J Cell Physiol. 1995;165(2):307–312. doi:10.1002/JCP.1041650211

63. Miller KJ, Thaloor D, Matteson S, Pavlath GK. Hepatocyte growth factor affects satellite cell activation and differentiation in regenerating skeletal muscle. Am J Physiol Cell Physiol. 2000;278(1). doi:10.1152/AJPCELL.2000.278.1.C174

64. Tatsumi R, Anderson JE, Nevoret CJ, Halevy O, Allen RE. HGF/SF is present in normal adult skeletal muscle and is capable of activating satellite cells. Dev Biol. 1998;194(1):114–128. doi:10.1006/DBIO.1997.8803

65. Cooper RN, Tajbakhsh S, Mouly V, Cossu G, Buckingham M, Butler-Browne GS. In vivo satellite cell activation via Myf5 and MyoD in regenerating mouse skeletal muscle. J Cell Sci. 1999;112 (Pt 17)(17):2895–2901. doi:10.1242/JCS.112.17.2895

66. Flamini V, Ghadiali RS, Antczak P, Rothwell A, Turnbull JE, Pisconti A. The Satellite Cell Niche Regulates the Balance between Myoblast Differentiation and Self-Renewal via p53. Stem Cell Reports. 2018;10(3):970–983. doi:10.1016/J.STEMCR.2018.01.007

67. Bentzinger CF, Wang YX, Rudnicki MA. Building Muscle: Molecular Regulation of Myogenesis. Cold Spring Harb Perspect Biol. 2012;4(2):a008342. doi:10.1101/CSHPERSPECT.A008342

68. Wang YX, Dumont NA, Rudnicki MA. Muscle stem cells at a glance. J Cell Sci. 2014;127(Pt 21):4543–4548. doi:10.1242/JCS.151209

69. Nguyen JH, Chung JD, Lynch GS, Ryall JG. The Microenvironment Is a Critical Regulator of Muscle Stem Cell Activation and Proliferation. Front Cell Dev Biol. 2019;7:254. doi:10.3389/fcell.2019.00254

70. Ruiz-Gómez M, Coutts N, Suster ML, Landgraf M, Bate M. myoblasts incompetent encodes a zinc finger transcription factor required to specify fusion-competent myoblasts in Drosophila. Development. 2002;129(1):133–141. doi:10.1242/DEV.129.1.133

71. Abmayr SM, Pavlath GK. Myoblast fusion: lessons from flies and mice. Development. 2012;139(4):641–656. doi:10.1242/DEV.068353

72. Isesele PO, Mazurak VC. Regulation of Skeletal Muscle Satellite Cell Differentiation by Omega-3 Polyunsaturated Fatty Acids: A Critical Review. Front Physiol. 2021;12:750. doi:10.3389/fphys.2021.682091

73. Arsic N, Zacchigna S, Zentilin L, et al. Vascular endothelial growth factor stimulates skeletal muscle regeneration in Vivo. Mol Ther. 2004;10(5):844–854. doi:10.1016/J.YMTHE.2004.08.007

74. Sonnet C, Lafuste P, Arnold L, et al. Human macrophages rescue myoblasts and myotubes from apoptosis through a set of adhesion molecular systems. J Cell Sci. 2006;119(Pt 12):2497–2507. doi:10.1242/JCS.02988

75. Cencetti F, Bernacchioni C, Tonelli F, Roberts E, Donati C, Bruni P. TGFβ1 evokes myoblast apoptotic response via a novel signaling pathway involving S1P4 transactivation upstream of Rho-kinase-2 activation. FASEB J. 2013;27(11):4532–4546. doi:10.1096/FJ.13-228528

76. Murphy MM, Lawson JA, Mathew SJ, Hutcheson DA, Kardon G. Satellite cells, connective tissue fibroblasts and their interactions are crucial for muscle regeneration. Development. 2011;138(17):3625–3637. doi:10.1242/dev.064162

77. Beanes SR, Dang C, Soo C, Ting K. Skin repair and scar formation: the central role of TGF-β. Expert Rev Mol Med. 2003;5(8):1–22. doi:10.1017/S1462399403005817

78. Chellini F, Tani A, Zecchi-Orlandini S, Sassoli C. Influence of platelet-rich and platelet-poor plasma on endogenous mechanisms of skeletal muscle repair/regeneration. Int J Mol Sci. 2019;20(3). doi:10.3390/ijms20030683

79. Dickinson RB, Guido S, Tranquillo RT. Biased cell migration of fibroblasts exhibiting contact guidance in oriented collagen gels. Ann Biomed Eng. 1994;22(4):342–356. doi:10.1007/BF02368241

80. Zou Y, Zhang RZ, Sabatelli P, Chu ML, Bönnemann CG. Muscle interstitial fibroblasts are the main source of collagen VI synthesis in skeletal muscle: implications for congenital muscular dystrophy types Ullrich and Bethlem. J Neuropathol Exp Neurol. 2008;67(2):144–154. doi:10.1097/NEN.0B013E3181634EF7

81. Sanderson RD, Fitch JM, Linsenmayer TR, Mayne R. Fibroblasts promote the formation of a continuous basal lamina during myogenesis in vitro. J Cell Biol. 1986;102(3):740. doi:10.1083/JCB.102.3.740

82. Yokoyama T, Sekiguchi K, Tanaka T, et al. Angiotensin II and mechanical stretch induce production of tumor necrosis factor in cardiac fibroblasts. Am J Physiol. 1999;276(6). doi:10.1152/AJPHEART.1999.276.6.H1968

83. Skutek M, Van Griensven M, Zeichen J, Brauer N, Bosch U. Cyclic mechanical stretching modulates secretion pattern of growth factors in human tendon fibroblasts. Eur J Appl Physiol. 2001;86(1):48–52. doi:10.1007/S004210100502

84. Lindner D, Zietsch C, Becher PM, et al. Differential expression of matrix metalloproteases in human fibroblasts with different origins. Biochem Res Int. Published online 2012. doi:10.1155/2012/875742

85. Newman AC, Nakatsu MN, Chou W, Gershon PD, Hughes CCW. The requirement for fibroblasts in angiogenesis: Fibroblast-derived matrix proteins are essential for endothelial cell lumen formation. Mol Biol Cell. 2011;22(20):3791–3800. doi:10.1091/mbc.E11-05-0393

86. Desmouliere A, Geinoz A, Gabbiani F, Gabbiani G. Transforming growth factor-beta 1 induces alpha-smooth muscle actin expression in granulation tissue myofibroblasts and in quiescent and growing cultured fibroblasts. J Cell Biol. 1993;122(1):103–111. doi:10.1083/JCB.122.1.103

87. Wipff PJ, Rifkin DB, Meister JJ, Hinz B. Myofibroblast contraction activates latent TGF-beta1 from the extracellular matrix. J Cell Biol. 2007;179(6):1311–1323. doi:10.1083/JCB.200704042

88. Petrov V V., Fagard RH, Lijnen PJ. Stimulation of Collagen Production by Transforming Growth Factor-β1 During Differentiation of Cardiac Fibroblasts to Myofibroblasts. Hypertension. 2002;39(2 I):258–263. doi:10.1161/HY0202.103268

89. Lemos DR, Babaeijandaghi F, Low M, et al. Nilotinib reduces muscle fibrosis in chronic muscle injury by promoting TNF-mediated apoptosis of fibro/adipogenic progenitors. Nat Med. 2015;21(7):786–794. doi:10.1038/NM.3869

90. Kim J, Lee J. Role of transforming growth factor-β in muscle damage and regeneration: focused on eccentric muscle contraction. J Exerc Rehabil. 2017;13(6):621. doi:10.12965/JER.1735072.536

91. Huey KA. Potential Roles of Vascular Endothelial Growth Factor during Skeletal Muscle Hypertrophy. Exerc Sport Sci Rev. 2018;46(3):195–202. doi:10.1249/JES.0000000000000152

92. Snijders T, Nederveen JP, McKay BR, et al. Satellite cells in human skeletal muscle plasticity. Front Physiol. 2015;6(OCT):283. doi:10.3389/fphys.2015.00283

93. Wickler SJ. Capillary supply of skeletal muscles from acclimatized white-footed mice Peromyscus. Am J Physiol. 1981;241(5):357–361. doi:10.1152/AJPREGU.1981.241.5.R357

94. Gehlert S, Theis C, Weber S, et al. Exercise-induced decline in the density of lyve-1-positive lymphatic vessels in human skeletal muscle. Lymphat Res Biol. 2010;8(3):165–173. doi:10.1089/LRB.2009.0035

95. Jacobsen NL, Norton CE, Shaw RL, Cornelison D, Segal S, Segal SS. Myofiber injury induces capillary disruption and regeneration of disorganized microvascular networks. bioRxiv. Published online August 4, 2021:2021.08.02.454805. doi:10.1101/2021.08.02.454805

96. Frey SP, Jansen H, Raschke MJ, Meffert RH, Ochman S. VEGF improves skeletal muscle regeneration after acute trauma and reconstruction of the limb in a rabbit model. Clin Orthop Relat Res. 2012;470(12):3607–3614. doi:10.1007/S11999-012-2456-7

97. Hardy D, Besnard A, Latil M, et al. Comparative Study of Injury Models for Studying Muscle Regeneration in Mice. PLoS One. 2016;11(1):e0147198. doi:10.1371/JOURNAL.PONE.0147198

98. Haas TL, Milkiewicz M, Davis SJ, et al. Matrix metalloproteinase activity is required for activity-induced angiogenesis in rat skeletal muscle. Am J Physiol Heart Circ Physiol. 2000;279(4). doi:10.1152/AJPHEART.2000.279.4.H1540

99. Hampton HR, Chtanova T. Lymphatic Migration of Immune Cells. Front Immunol. 2019;10(MAY):1168. doi:10.3389/FIMMU.2019.01168

100. ExternalPotential Plugin — CC3D Reference Manual 4.4.1 documentation. Accessed January 4, 2024. https://compucell3dreferencemanual.readthedocs.io/en/latest/external_potential_plugin.ht ml

101. Filion RJ, Popel AS. Intracoronary administration of FGF-2: A computational model of myocardial deposition and retention. Am J Physiol - Hear Circ Physiol. 2005;288(1 57-1):263–279. doi:10.1152/ajpheart.00205.2004

102. Ochoa O, Sun D, Reyes-Reyna SM, et al. Delayed angiogenesis and VEGF production in CCR2-/-mice during impaired skeletal muscle regeneration. Am J Physiol - Regul Integr Comp Physiol. 2007;293(2):651–661. doi:10.1152/ajpregu.00069.2007

103. Pratt SJP, Shah SB, Ward CW, Kerr JP, Stains JP, Lovering RM. Recovery of altered neuromuscular junction morphology and muscle function in mdx mice after injury. Cell Mol Life Sci. 2014;72(1):153–164. doi:10.1007/s00018-014-1663-7

104. You JS, Barai P, Chen J. Sex differences in skeletal muscle size, function, and myosin heavy chain isoform expression during post-injury regeneration in mice. Physiol Rep. 2023;11(16). doi:10.14814/PHY2.15791

105. Marino S, Hogue IB, Ray CJ, Kirschner DE. A methodology for performing global uncertainty and sensitivity analysis in systems biology. J Theor Biol. 2008;254(1):178–196. doi:10.1016/J.JTBI.2008.04.011

106. Segovia-Juarez JL, Ganguli S, Kirschner D. Identifying control mechanisms of granuloma formation during M. tuberculosis infection using an agent-based model. J Theor Biol. 2004;231(3):357–376. doi:10.1016/J.JTBI.2004.06.031

107. Wang X, Zhao W, Ransohoff RM, Zhou L. Infiltrating macrophages are broadly activated at the early stage to support acute skeletal muscle injury repair. J Neuroimmunol. 2018;317:55. doi:10.1016/J.JNEUROIM.2018.01.004

108. Nguyen MH, Cheng M, Koh TJ. Impaired Muscle Regeneration in Ob/ob and Db/db Mice. Sci World J. 2011;11:1525. doi:10.1100/TSW.2011.137

109. Teixeira CFP, Zamunér SR, Zuliani JP, et al. Neutrophils do not contribute to local tissue damage, but play a key role in skeletal muscle regeneration, in mice injected with Bothrops asper snake venom. Muscle Nerve. 2003;28(4):449–459. doi:10.1002/MUS.10453

110. Liu X, Liu Y, Zhao L, Zeng Z, Xiao W, Chen P. Macrophage depletion impairs skeletal muscle regeneration: The roles of regulatory factors for muscle regeneration. Cell Biol Int. 2017;41(3):228–238. doi:10.1002/CBIN.10705

111. Hardy D, Fefeu M, Besnard A, et al. Defective angiogenesis in CXCL12 mutant mice impairs skeletal muscle regeneration. Skelet Muscle. 2019;9(1):1–15. doi:10.1186/s13395-019-0210-5

112. Cantaert T, Baeten D, Tak PP, van Baarsen LGM. Type I IFN and TNFα cross-regulation in immune-mediated inflammatory disease: Basic concepts and clinical relevance. Arthritis Res Ther. 2010;12(5):1–10. doi:10.1186/AR3150/TABLES/2

113. Kunze KN, Hannon CP, Fialkoff JD, Frank RM, Cole BJ. Platelet-rich plasma for muscle injuries: A systematic review of the basic science literature. World J Orthop. 2019;10(7):278–291. doi:10.5312/wjo.v10.i7.278

114. Choi W, Lee J, Lee J, Lee SH, Kim S. Hepatocyte growth factor regulates macrophage transition to the M2 phenotype and promotes murine skeletal muscle regeneration. Front Physiol. 2019;10(JUL). doi:10.3389/fphys.2019.00914

115. Lu H, Huang D, Ransohoff RM, Zhou L. Acute skeletal muscle injury: CCL2 expression by both monocytes and injured muscle is required for repair. FASEB J. 2011;25(10):3344–3355. doi:10.1096/FJ.10-178939

116. Girardi F, Taleb A, Ebrahimi M, et al. TGFβ signaling curbs cell fusion and muscle regeneration. Nat Commun. 2021;12(1). doi:10.1038/s41467-020-20289-8

117. Zimowska M, Olszynski KH, Swierczynska M, Streminska W, Ciemerych MA. Decrease of MMP-9 activity improves soleus muscle regeneration. Tissue Eng - Part A. 2012;18(11-12):1183–1192. doi:10.1089/ten.tea.2011.0459

118. Dziki JL, Velayutham M, Hussey GS, Turnquist HR. Cytokine networks in immune-mediated muscle regeneration. J Immunol Regen Med. 2018;1:32–44. doi:10.1016/j.regen.2018.03.001

119. Fujita R, Kawano F, Ohira T, et al. Anti-interleukin-6 receptor antibody (MR16-1) promotes muscle regeneration via modulation of gene expressions in infiltrated macrophages. Biochim Biophys Acta - Gen Subj. 2014;1840(10):3170–3180. doi:10.1016/j.bbagen.2014.01.014

120. Ismail AA, Shaker BT, Bajou K. The plasminogen–activator plasmin system in physiological and pathophysiological angiogenesis. Int J Mol Sci. 2022;23(1). doi:10.3390/ijms23010337

121. Alsousou J, Ali A, Willett K, Harrison P. The role of platelet-rich plasma in tissue regeneration. Platelets. 2013;24(3):173–182. doi:10.3109/09537104.2012.684730

122. Van De Kamp J, Jahnen-Dechent W, Rath B, Knuechel R, Neuss S. Hepatocyte Growth Factor-Loaded Biomaterials for Mesenchymal Stem Cell Recruitment. Stem Cells Int. 2013;2013. doi:10.1155/2013/892065

123. Akhurst RJ. TGF-β antagonists: Why suppress a tumor suppressor? J Clin Invest. 2002;109(12):1533. doi:10.1172/JCI15970

124. Li H, Mittal A, Makonchuk DY, Bhatnagar S, Kumar A. Matrix metalloproteinase-9 inhibition ameliorates pathogenesis and improves skeletal muscle regeneration in muscular dystrophy. Hum Mol Genet. 2009;18(14):2584. doi:10.1093/HMG/DDP191

125. Lin CC, Boyer PD, Aimetti AA, Anseth KS. Regulating MCP-1 Diffusion in Affinity Hydrogels for Enhancing Immuno-isolation. J Control Release. 2010;142(3):384. doi:10.1016/J.JCONREL.2009.11.022

126. Marino S, Hult C, Wolberg P, Linderman JJ, Kirschner DE. The Role of Dimensionality in Understanding Granuloma Formation. *Comput (Basel*, Switzerland). 2018;6(4). doi:10.3390/COMPUTATION6040058

127. Sego TJ, Kasacheuski U, Hauersperger D, Tovar A, Moldovan NI. A heuristic computational model of basic cellular processes and oxygenation during spheroid-dependent biofabrication. Biofabrication. 2017;9(2):024104. doi:10.1088/1758-5090/AA6ED4

128. Haizlip KM, Harrison BC, Leinwand LA. Sex-based differences in skeletal muscle kinetics and fiber-type composition. Physiology. 2015;30(1):30–39. doi:10.1152/physiol.00024.2014

129. Knewtson KE, Ohl NR, Robinson JL. Estrogen Signaling Dictates Musculoskeletal Stem Cell Behavior: Sex Differences in Tissue Repair. Tissue Eng - Part B Rev. 2022;28(4):789–812. doi:10.1089/ten.teb.2021.0094

130. Liu S, Kostek M, Omstead K. Sex based differences in muscle regeneration. Physiology. 2023;38(S1). doi:10.1152/physiol.2023.38.s1.5729941

131. Enns DL, Tiidus PM. The influence of estrogen on skeletal muscle: Sex matters. Sport Med. 2010;40(1):41–58. doi:10.2165/11319760-000000000-00000

132. Zhao W, Zhao H, Li M, Huang C. Microfluidic devices for neutrophil chemotaxis studies. J Transl Med. 2020;18(1):1–19. doi:10.1186/s12967-020-02335-7

133. Heit B, Liu L, Colarusso P, Puri KD, Kubes P. PI3K accelerates, but is not required for, neutrophil chemotaxis to fMLP. J Cell Sci. 2008;121(2):205–214. doi:10.1242/JCS.020412

134. Martin KS, Kegelman CD, Virgilio KM, et al. In Silico and In Vivo Experiments Reveal M-CSF Injections Accelerate Regeneration Following Muscle Laceration. Ann Biomed Eng. 45. doi:10.1007/s10439-016-1707-2

135. Bos E van den, Walbaum S, Horsthemke M, Bachg AC, Hanley PJ. Time-lapse Imaging of Mouse Macrophage Chemotaxis. JoVE (Journal Vis Exp. 2020;2020(158):e60750. doi:10.3791/60750

136. Corliss BA, Azimi MS, Munson JM, Peirce SM, Murfee WL. Macrophages: An Inflammatory Link between Angiogenesis and Lymphangiogenesis. Microcirculation. 2016;23(2):95. doi:10.1111/MICC.12259

137. Newby AC. Metalloproteinase expression in monocytes and macrophages and its relationship to atherosclerotic plaque instability. Arterioscler Thromb Vasc Biol. 2008;28(12):2108–2114. doi:10.1161/ATVBAHA.108.173898

138. Lu HL, Huang XY, Luo YF, Tan WP, Chen PF, Guo YB. Activation of M1 macrophages plays a critical role in the initiation of acute lung injury. Biosci Rep. 2018;38(2):20171555. doi:10.1042/BSR20171555

139. Cui K, Ardell CL, Podolnikova NP, Yakubenko VP. Distinct migratory properties of M1, M2, and resident macrophages are regulated by αdβ2and αmβ2integrin-mediated adhesion. Front Immunol. 2018;9(NOV):2650. doi:10.3389/fimmu.2018.02650

140. da Silva MD, Bobinski F, Sato KL, Kolker SJ, Sluka KA, Santos ARS. IL-10 Cytokine Released from M2 Macrophages Is Crucial for Analgesic and Anti-inflammatory Effects of Acupuncture in a Model of Inflammatory Muscle Pain. Mol Neurobiol. 2015;51(1):19. doi:10.1007/S12035-014-8790-X

141. Moncayo AR. Poisson Convergence and Family Trees. Ann Probab. 2007;3(6). doi:10.1214/aop/1176996235

142. Xiaoping C, Yong L. Role of matrix metalloproteinases in skeletal muscle: migration, differentiation, regeneration and fibrosis. Cell Adh Migr. 2009;3(4). doi:10.4161/CAM.3.4.9338

143. Terrente Y, El Fahime E, Caron NJ, et al. Tumor necrosis factor-alpha (TNF-alpha) stimulates chemotactic response in mouse myogenic cells. Cell Transplant. 2003;12(1):91–100. doi:10.3727/000000003783985115

144. Bakkar N, Wang J, Ladner KJ, et al. IKK/NF-κB regulates skeletal myogenesis via a signaling switch to inhibit differentiation and promote mitochondrial biogenesis. J Cell Biol. 2008;180(4):787. doi:10.1083/JCB.200707179

145. Saclier M, Yacoub-Youssef H, Mackey AL, et al. Differentially activated macrophages orchestrate myogenic precursor cell fate during human skeletal muscle regeneration. Stem Cells. 2013;31(2):384–396. doi:10.1002/STEM.1288

146. Perandini LA, Chimin P, Lutkemeyer D da S, Câmara NOS. Chronic inflammation in skeletal muscle impairs satellite cells function during regeneration: can physical exercise restore the satellite cell niche? FEBS J. 2018;285(11):1973–1984. doi:10.1111/FEBS.14417

147. Gal-Levi R, Leshem Y, Aoki S, Nakamura T, Halevy O. Hepatocyte growth factor plays a dual role in regulating skeletal muscle satellite cell proliferation and differentiation. Biochim Biophys Acta - Mol Cell Res. 1998;1402(1):39–51. doi:10.1016/S0167-4889(97)00124-9

148. Ten Broek RW, Grefte S, Von Den Hoff JW. Regulatory factors and cell populations involved in skeletal muscle regeneration. J Cell Physiol. 2010;224(1):7–16. doi:10.1002/JCP.22127

149. Kuang S, Kuroda K, Le Grand F, Rudnicki MA. Asymmetric self-renewal and commitment of satellite stem cells in muscle. Cell. 2007;129(5):999–1010. doi:10.1016/J.CELL.2007.03.044

150. Yennek S, Burute M, Théry M, Tajbakhsh S. Cell adhesion geometry regulates non-random DNA segregation and asymmetric cell fates in mouse skeletal muscle stem cells. Cell Rep. 2014;7(4):961–970. doi:10.1016/J.CELREP.2014.04.016

151. Siegel AL, Kuhlmann PK, Cornelison DDW. Muscle satellite cell proliferation and association: new insights from myofiber time-lapse imaging. Skelet Muscle. 2011;1(1). doi:10.1186/2044-5040-1-7

152. Rocheteau P, Gayraud-Morel B, Siegl-Cachedenier I, Blasco MA, Tajbakhsh S. A subpopulation of adult skeletal muscle stem cells retains all template DNA strands after cell division. Cell. 2012;148(1-2):112–125. doi:10.1016/J.CELL.2011.11.049

153. Otto A, Collins-Hooper H, Patel A, Dash PR, Patel K. Adult skeletal muscle stem cell migration is mediated by a blebbing/amoeboid mechanism. Rejuvenation Res. 2011;14(3):249–260. doi:10.1089/rej.2010.1151

154. Gibb AA, Lazaropoulos MP, Elrod JW. Myofibroblasts and Fibrosis. Circ Res. 2020;127(3):427-447. doi:10.1161/CIRCRESAHA.120.316958

155. Alberts B, Johnson A, Lewis J, Raff M, Roberts K, Walter P. Extracellular Control of Cell Division, Cell Growth, and Apoptosis. Published online 2002. Accessed March 14, 2023. https://www.ncbi.nlm.nih.gov/books/NBK26877/

156. Cornwell KG, Downing BR, Pins GD. Characterizing fibroblast migration on discrete collagen threads for applications in tissue regeneration. J Biomed Mater Res Part A. 2004;71A(1):55-62. doi:10.1002/JBM.A.30132

157. Qutub A, Gabhann F, Karagiannis E, Vempati P, Popel A. Multiscale models of angiogenesis. In: IEEE Engineering in Medicine and Biology Magazine. Vol 28. NIH Public Access; 2009:14–31. doi:10.1109/MEMB.2009.931791

158. Tigner A, Ibrahim SA, Murray I. Histology, White Blood Cell. StatPearls. Published online November 19, 2021. Accessed May 16, 2022. https://www.ncbi.nlm.nih.gov/books/NBK563148/

159. Garcia SM, Tamaki S, Lee S, et al. Stem Cell Reports Resource High-Yield Purification, Preservation, and Serial Transplantation of Human Satellite Cells. Published online 2018. doi:10.1016/j.stemcr.2018.01.022

160. Krombach F, Münzing S, Allmeling AM, Gerlach JT, Behr J, Dörger M. Cell size of alveolar macrophages: an interspecies comparison. Environ Health Perspect. 1997;105(Suppl 5):1261. doi:10.1289/EHP.97105S51261

161. Downey GP, Doherty DE, Schwab B, Elson EL, Henson PM, Worthen GS. Retention of leukocytes in capillaries: role of cell size and deformability. 101152/jappl19906951767. 1990;69(5):1767–1778. doi:10.1152/JAPPL.1990.69.5.1767

162. Freitas RJ. Nanomedicine: Basic Capabilities. Vol 1. Landes Bioscience; 1999. Accessed May 16, 2022. http://www.nanomedicine.com/NMI/8.5.1.htm

163. Deng B, Wehling-Henricks M, Villalta SA, Wang Y, Tidball JG. IL-10 Triggers Changes in Macrophage Phenotype That Promote Muscle Growth and Regeneration. J Immunol. 2012;189(7):3669–3680. doi:10.4049/jimmunol.1103180

164. Raimondo TM, Mooney DJ. Functional muscle recovery with nanoparticle-directed M2 macrophage polarization in mice. Proc Natl Acad Sci U S A. 2018;115(42):10648–10653. doi:10.1073/pnas.1806908115

165. Chen SE, Gerken E, Zhang Y, et al. Role of TNF-α signaling in regeneration of cardiotoxin-injured muscle. Am J Physiol Cell Physiol. 2005;289(5):C1179. doi:10.1152/AJPCELL.00062.2005

